# *Oryza* genera-specific novel Histone H4 variant predisposes H4 Lysine5 Acetylation marks to modulate salt stress responses

**DOI:** 10.1101/2023.07.31.551207

**Authors:** Hari Sundar G Vivek, Paula Sotelo-Parrilla, Steffi Raju, Shaileshanand Jha, Anjitha Gireesh, Fabian Gut, K.R. Vinothkumar, Frédéric Berger, A. Arockia Jeyaprakash, P.V. Shivaprasad

## Abstract

Paralogous variants of canonical histones guide accessibility to DNA and function as additional layers of genome regulation. Across eukaryotes, the occurrence, mechanism of action and functional significance of several variants of core histones are well known except that of histone H4. Here we show that a novel variant of H4 (H4.V), expressing tissue-specifically among members of *Oryza* genera, mediates specific epigenetic changes contributing majorly to salt tolerance. H4.V was incorporated to specific chromosomal locations where it blocked deposition of active histone marks. Under salt stress, large scale re-distribution of H4.V enabled incorporation of stress dependent histone H4 Lysine5 Acetylation (H4K5Ac) marks. Mis-expression of H4.V led to defects at morphological level especially in reproductive tissues, and in mounting stress responses. H4.V mediated these alterations by condensing chromatin at specific genomic regions as seen with cryo-EM structure of reconstituted H4.V containing nucleosomes. These results not only uncovered the presence of a H4 variant in plants, but also a novel chromatin regulation of stress responses that might have contributed to success of semi-aquatic *Oryza* members under variable water-limiting conditions.

**One-line summary:** Histone H4 variant predisposes chromatin for stress responses

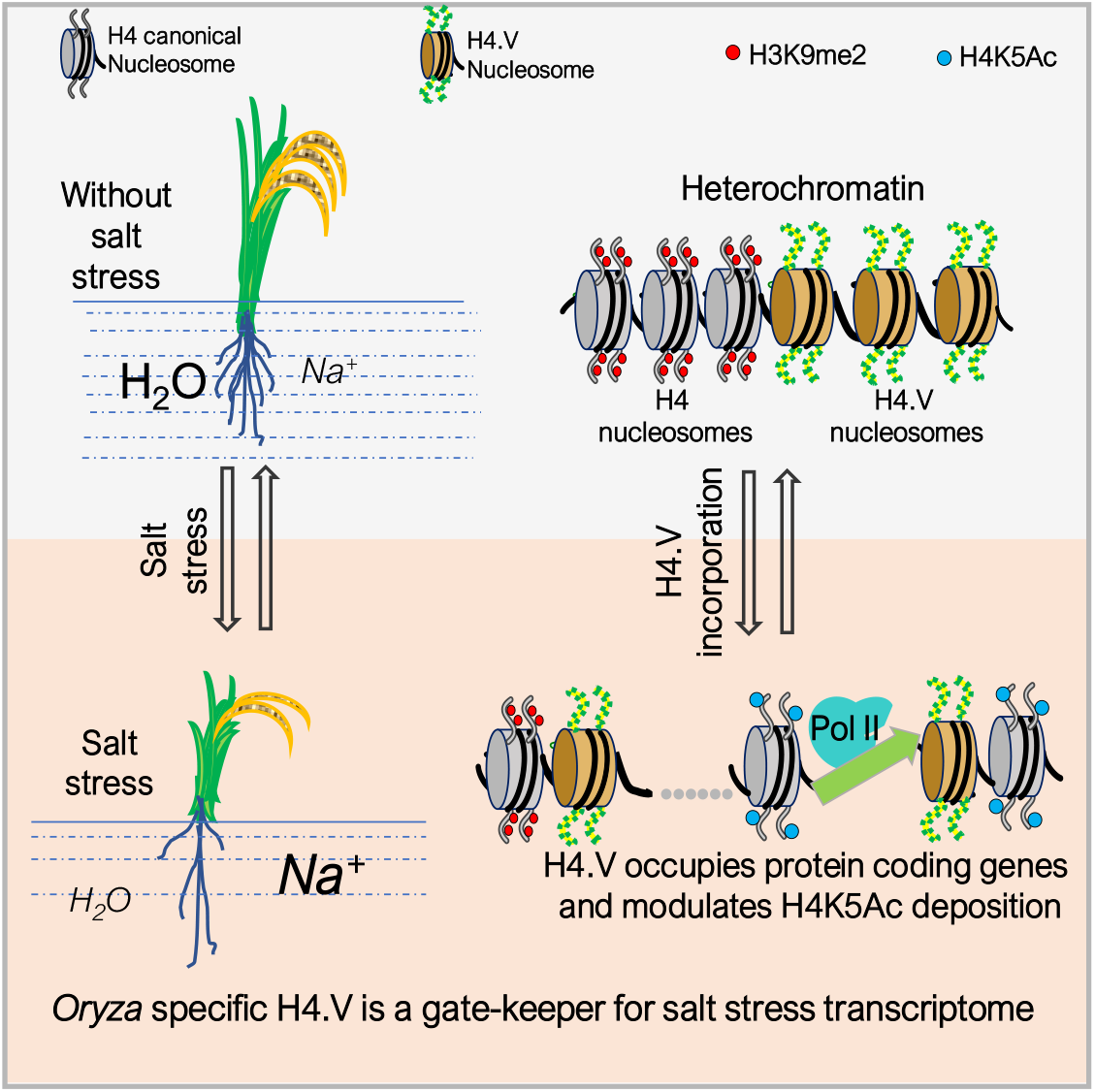

## Introduction

Histones are modular proteins conserved across eukaryotes aiding in packaging the nuclear DNA as chromatin (Kornberg, 1974). Excluding archaea and few instances of eubacteria (Hocher *et al*, 2023; Henneman *et al*, 2018; Mattiroli *et al*, 2017), roughly 147 bp DNA is wrapped around an octamer of two units of histones H2A, H2B, H3 and H4, forming nucleosomes in eukaryotes (Luger *et al*, 1997). In addition to DNA modifications, nucleosomal histones undergo several post-translational modifications (PTMs), predominantly at the tails, some of which reflect transcriptional activity and chromatin architecture (Demetriadou *et al*, 2020; Bannister & Kouzarides, 2011). Several paralogs of H2A, H2B and H3 histones, named histone variants, have been identified across eukaryotes that are expressed and incorporated into the chromatin independent of cell cycle (Talbert & Henikoff, 2010). H3 and H2A.Z variants are distinguished from their counterparts by specific histone chaperone complexes that assist their incorporation in chromatin and often perform essential roles (Otero *et al*, 2014; Henikoff, 2008).

By virtue of their specific residue modifications and variations, it has been shown that H2A variants confer unique properties to the nucleosomes thus regulating chromatin (Osakabe *et al*, 2018; Talbert & Henikoff, 2021). In addition to altering the chromatin properties, specific histone variants facilitate essential cellular processes such as chromosome segregation during cell division (CENH3), DNA damage responses (H2A.X), and gametic inheritance of epigenetic states (H3.10) (Ravi & Chan, 2010; Schmücker *et al*, 2021; Lorković *et al*, 2017; Buttress *et al*, 2022; Jiang *et al*, 2020; Jamge *et al*, 2023). H2A.W was identified as a plant specific histone variant that functions in efficient silencing of transposons at the heterochromatin, acting as a determinant of several repressive chromatin states (Yelagandula *et al*, 2014; Osakabe *et al*, 2021; Jamge *et al*, 2023).

Multiple functionally relevant H2A, H2B and H3 variants have been described among plants and animals (Talbert & Henikoff, 2017). However, only a very few variants in H4 has been described and all were in parasites (Alsford & Horn, 2004) with the exception of H4 variant (H4G) specific to the *Hominidae* family has been characterised as a regulator of rDNA expression in specific cancer cells (Long *et al*, 2019; Pang *et al*, 2020). In contrast, histone H4 sequences are highly conserved and variants of histone H4 have not been functionally characterised in plants. Evolution of monophyletic clade-specific histone variants having very specific functions have been documented in several cases (Jiang *et al*, 2020; Chang *et al*, 2023; Raman *et al*, 2022). In such scenarios, repurposing of the canonical histone chaperones enabled incorporation of the variants (Long *et al*, 2019), indicating that the functional outcome largely dependent on variants than their chaperones.

Plant responses are unique to different types of stresses and are distinct in different species (Zhang *et al*, 2022; Pandey *et al*, 2015). Tuned responses to stress is important for tolerance and plants have evolved several layers of regulatory modules to counteract aberrant activation of stress responsive genes (Zhu, 2016; Bartels & Sunkar, 2005). Chromatin level changes upon stress was identified as a powerful module of stress responses as several genes were regulated rapidly (Lämke & Bäurle, 2017; Probst & Mittelsten Scheid, 2015). Since plants can produce gametes from any tissue and do not segregate gametic cell lineage, the stress induced epigenetic changes gets effectively transmitted across mitosis and meiosis, thereby promoting transgenerational memory of the stress (Mladenov *et al*, 2021). Histone deacetylases and other modifiers were implicated in modulating global stress responses via genome-wide changes in reprogramming of histone modifications (Luo *et al*, 2017; Li *et al*, 2021). These modifiers regulated histone marks such as H4K5Ac or H3K4me, thereby regulating stress responsive TFs (He *et al*, 2022; Xu *et al*, 2022; He *et al*, 2020). Histone variants were also found to initiate signaling during abiotic stress responses (Nunez-Vazquez *et al*, 2022). Importance of H2A.Z has been well-documented during heat and other stresses, where H2A.Z containing nucleosomes poise transcription for stress tolerance and responsive genes (Berriri *et al*, 2016; March-Díaz *et al*, 2008; Sura *et al*, 2017; Coleman-Derr & Zilberman, 2012; Kumar & Wigge, 2010). However, roles of other histone variants or mechanism by which the histone variants modulate the chromatin to facilitate stress responses are not understood. Most of the above studies were carried out in a seasonal model plant *Arabidopsis* having small homogenous compartmentalized genome, while the diversity, complexity, properties and regulation of chromatin, among plants with larger genomes such as crops, are manifold heterogenous and often species-specific.

In this work, we identified and functionally characterized a novel *Oryza* genera specific histone H4 variant that exhibited sequence conservation within the genera. This is the first report to functionally characterize a variant of H4 in plants. Using molecular, genetic, and genomic approaches, we established the functional role of the H4.V in rice. Analyses of cryo-EM structures and biochemical analyses of the H4.V and H4 containing rice nucleosomes revealed unique properties of the H4.V that can aid in chromatin regulation. Genetic studies with mis-expression lines including mutants of the gene indicated its surprising and unique roles in synergistic modulation of H4K5Ac chromatin marks under salt stress conditions. These results indicate that the adaptation of semi-aquatic *Oryza* members to changing environments was likely mediated through a significant contribution from this novel H4 variant.

## Results

### Novel *Oryza* genera specific histone H4 variant predominantly occupied intergenic loci

Most eukaryotic genomes comprise of multiple histone genes, occurring in clusters or otherwise, that generate histones upon entering S-phase of cell-cycle, feeding to the chromatin assembly factors (Armstrong & Spencer, 2021). Likewise, rice genome comprises ten histone H4 genes of which nine encode histones identical to canonical H4 (named H4 throughout) except for the gene locus LOC_Os05g38760 (MSU gene model) (Appendix Fig S1A). This gene locus encoded a previously uncharacterized histone H4 like protein that we named as H4.V. Sequence alignment indicated that H4.V resembled H4 in length (103 amino acids) with similar histone fold domains albeit with minor modifications (Fig 1A). Majority of the variations in sequence were confined to the N-terminal tail, with the first 27 and 55 residues encompassing 50% and 70% of all the variations respectively (Fig 1B). Particularly, the first 20 residues had the most disfavoured amino acid substitutions on a BLOSUM62 substitution scale (Fig 1A). H4.V showed 72.8% sequence identity to H4 that was largely conserved across plants and metazoans (Fig 1C, Appendix Table S1). H4.V was almost identical among ancient rice species - *O. nivara, O. barthii, O. rufipogon,* and modern *japonica* and *indica* subspecies of *O. sativa* (Appendix Fig S1B). *H4.V* was present in all the 3001 sequenced rice (Appendix Fig S1C) but similar sequences were absent in other genera indicating that H4.V is likely *Oryza* specific.

**Figure 1.**
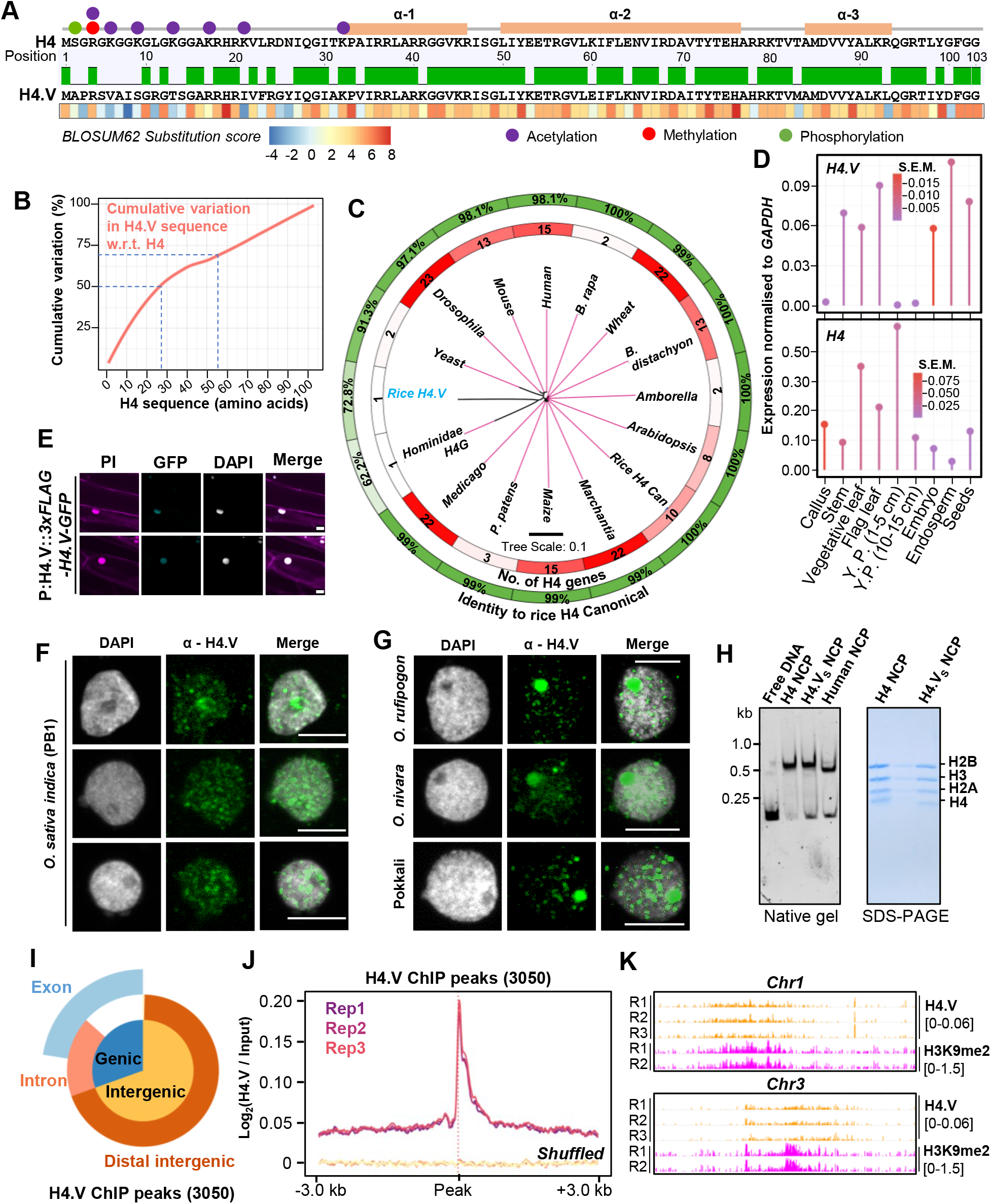
Rice specific histone H4.V is different from H4. (A) Pairwise sequence alignment of H4 and H4.V. Line diagram shows histone-fold domains (oran boxes) and H4 residues that attract PTMs (colour coded). Heatmap with BLOSUM62 score substitutions observed. (B) Cumulative variations plot showing positional distribution of v protein length from N-terminal end (x-axis). Dotted lines represent the intercep variation. (C) Phylogenetic tree with protein sequence variations of H4 ac number of H4 encoding genes; outer track-% identity w.r.t. rice H4. T plots showing RT-qPCR estimation of H4 and H4.V expression represent standard error of the mean (S.E.M.). Y.P.: seedling roots expressing 3xFLAG-H4.V-GFP driv (F), *Oryza* species and pokkali (G) nuclei s reconstituted H4.V_S_ and H4 containing NCPs on a native gel (6% acrylamide) and Coomassie brilliant blue (CBB) stained SDS-PAGE gel (15% acrylamide). Human NCPs were used as control. (I) Vennpie chart showing proportion of genic and intergenic regions (beyond +/- 3 kb of genes) overlapped by H4.V peaks. (J) Metagene plots showing enrichment of H4.V ChIP-seq signal in 3 biological replicates over H4.V peaks. Enrichment at the shuffled set of loci served as control. (K) Chromosome-wide genome browser screenshots showing ChIP enrichment (square brackets) of H4.V and H3K9me2 marks. (E-G) DAPI tained DNA; propidium iodide (PI) stained DNA and cell walls. Scale: 5μm.

*H4.V* was cloned from cDNA and its sequence matched the coding sequence in the reference genome (Appendix Fig S1D). Expression profiling revealed that *H4.V* expressed in a tissue-specific manner and overall transcript levels of *H4.V* were lower than *H4* (Fig 1D). When compared to *H4*, *H4.V* was predominantly expressed in embryos, endosperm and seeds and was absent in young panicles. GFP tagged H4.V fusion protein, driven by its native promoter, was localised to the nucleus in rice root cells (Fig 1E). In order to demonstrate that H4.V is a true variant capable of incorporation into the chromatin and to study its occupancy, we raised specific polyclonal antibodies (methods) that did not cross react with H4 (Appendix Fig S2A-D). Immunofluorescence (IFL) studies with this antibody revealed that H4.V is expressed and deposited in the chromatin of *indica* rice, its wild relatives including *O. nivara, O. rufipogon* and the locally cultivated landrace pokkali, suggesting the conservation of the variant among *Oryza* genera members (Figs 1F and G). To test if H4.V is incorporated into the nucleosomes, we purified rice histones H2A, H2B, H3, H4.V and H4 recombinantly in *E. coli* and performed gradual native buffer dialysis to enable refolding into octamers (Methods). The histone octamers were insoluble when folded *in vitro* with H4.V but were soluble when the N-terminal end (53 amino acids, accounting for most of the variation) of H4.V was fused with C-terminal 50 residues of H4 (named H4.V_S_ throughout) (Appendix Fig S3A-C). With recombinant H4.V_S_ or H4 and H2A, H2B and H3 from rice and Widom 601 nucleosome positioning DNA sequence (NPS) (Lowary & Widom, 1998), we successfully reconstituted nucleosome core particles (NCPs) (Methods, Appendix Table S5). NCPs of H4.V_S_ and H4 exhibited similar mobility shift profiles in gel shift assays and similar dimensional decay properties in dynamic light scattering (DLS) assays, suggesting that H4.V can be incorporated into the nucleosomes *in vitro* and matches the profile of characterized human nucleosomes (Fig 1H and Appendix Figs S3D and E).

Using chromatin immunoprecipitation followed by deep sequencing (ChIP-seq) with our specific antibody, binding sites of H4.V were characterized (Appendix Dataset S1, three biological replicates). We identified roughly 3050 high confidence peaks of H4.V enrichment that represented majorly intergenic regions (Fig 1I and J). We also generated T-DNA free Cas9 mediated knockout lines of H4.V (*h4.v KO*) (Appendix Fig S4) that showed reduced ChIP enrichment at the H4.V peaks in comparison to WT plants (Appendix Fig S2C). Also, *h4.v KO* nuclei did not show H4.V signals in IFL microscopy, suggesting specificity of the H4.V antibody (Appendix Fig S2B). H4.V peaks predominantly occupied pericentromeres enriched with repressive H3K9me2 marks (Fig 1K). Taken together, expression, *in vitro* and *in planta* incorporation of this previously uncharacterized *Oryza* genera specific histone H4 variant at specific loci suggested its role in chromatin regulation.

### H4.V promoted structurally condensed and less-stable nucleosomes

In order to investigate the differences brought about by the residue variations to the nucleosomes, we performed single particle cryo-EM analysis of nucleosomes reconstituted using recombinant histones with either canonical H4 or H4.V_S_ (Methods and Appendix Table S6). Processing of the datasets using the standard CryoSPARC workflow yielded three-dimensional maps for H4 and H4.V_S_ containing nucleosomes at 3.6 Å resolution (Appendix Fig S5). The overall structure of the rice H4 nucleosome was similar to the human nucleosome structure (PDB: 7XD1) with key exceptions. About 20 bp of DNA in NCP entry/exit positions in H4 nucleosome was not resolved (from about Super Helical Loop +5) suggesting the presence of an asymmetrically flexible DNA entry/exit regions (Fig 2A). Consistent with this, N-terminal αN helix of H3 which directly binds the entry/exit DNA and the H2A C-terminal region in the human nucleosome was also not resolved at this site in the H4 NCP. Interestingly, when the structures of rice H4.V_S_ NCP and rice canonical NCP were compared, unlike H4 NCP, both entry/exit DNA regions of H4.Vs NCP were well resolved. In correlation with the stabilised entry/exit DNA regions, both H3 αN helix and the H2A C-terminal tail were well resolved in the H4.V_S_ NCP. Interestingly, H4.V has an Alanine to Valine substitution at position 34 which appears to stabilize the H3 loop connecting the H3 αN helix and α1 helix (Fig 2B).

**Figure 2.**
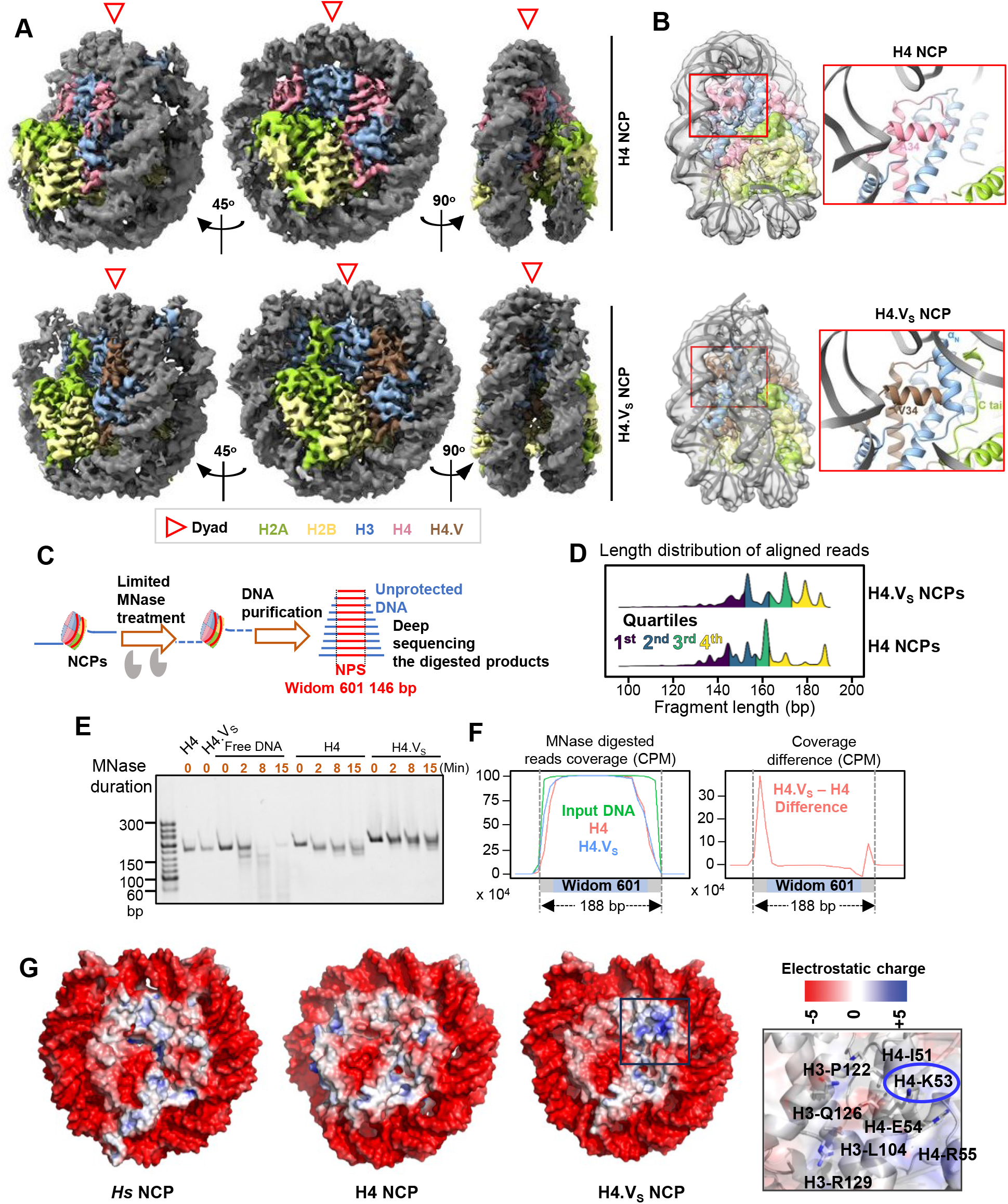
H4.V nucleosomes are structurally distinct to H4 nucleosomes. (A) The cryo-EM density map of H4 NCP and H4.Vs NCP as visualised by Chimera X. Unlike the H4.Vs NCP, in H4 NCP about 20 bp of one of the entry/exit DNA regions were not resolved highlighting the contribution of H4.Vs in stabilising the entry/exit DNA regions. (B) Close-up views of the H4 amino acid variation at position 34, which likely contributed to the stabilisation of H3 αN Helix, H2A C-terminal tail and DNA entry/exit region. (C) Schematic of the limited MNase digestion of the NCPs with extended Widom 601 sequence (NPS in red, extended DNA in blue). (D) Length distribution of purified DNA fragments after limited MNase digestion. Quartiles of length are coloured. (E) Representative gel showing the DNA size profiles of the MNase digestion over time course. (F) Coverage plots depicting density of reads over the extended Widom 601 NPS (left) and the difference in coverage plot between H4.V_S_ and H4 NCPs (right). (G) The electrostatic surface potential of human NCP (PDB:7XD1), H4 NCP and H4.Vs NCP. H4 NCP possesses an acidic patch that extends to H3/H4 when compared to human NCP where acidic patch is confined to H2A/H2B. Inset plot showing H4.V_S_ NCP with K53 residue highlighted that contributed to the charge variation.

In order to test if this structural feature of H4.V contributes to chromatin properties, we generated nucleosomes with H4.V_S_ and H4 octamers and Widom 601 NPS (146 bp) flanked by 21 bp free DNA on either side. NCPs were partially digested using micrococcal nuclease (MNase) which allows estimation of accessibility of free DNA, and the undigested DNA from the reaction mix was deep sequenced (Fig 2C). It was evident from the size distribution of DNA fragments that H4.V_S_ NCPs accumulated longer fragments, suggesting closer interaction of the DNA with the H4.V_S_ octamers when compared to H4 NCPs (Figs 2D,E). Further, alignment of the protected DNA fragments to the NPS revealed that the free DNA located at the termini of NPS was protected against MNase activity (Fig 2F). Thus the stabilization of entry/exit DNA regions observed in the H4.Vs NCP (Figs 2A,B) and altered electrostatic surface charge (Fig 2G) are consistent with our biochemical finding that the H4.Vs NCP is relatively more resistant to MNase digestion as compared to canonical NCP.

In order to further probe implications of variations in the H4.V N-terminal tail, we assayed several biochemical properties of the H4.V_S_ and H4 NCPs. Firstly, histone octamers (without DNA) prepared using H4.V_S_ were hydro-dynamically smaller than H4 containing octamers as they eluted later through a 24 ml gel filtration column (Fig 3A). The same results were observed prominently in a 120 ml size exclusion column signifying the variation of compaction between the H4.V_S_ and H4 containing histone octamers (Fig 3B). This observation of compact H4.V_S_ octamers corroborated the cryo-EM observations of stabilisation in H4.V_S_ NCP (Figs 2A,B).

**Figure 3.**
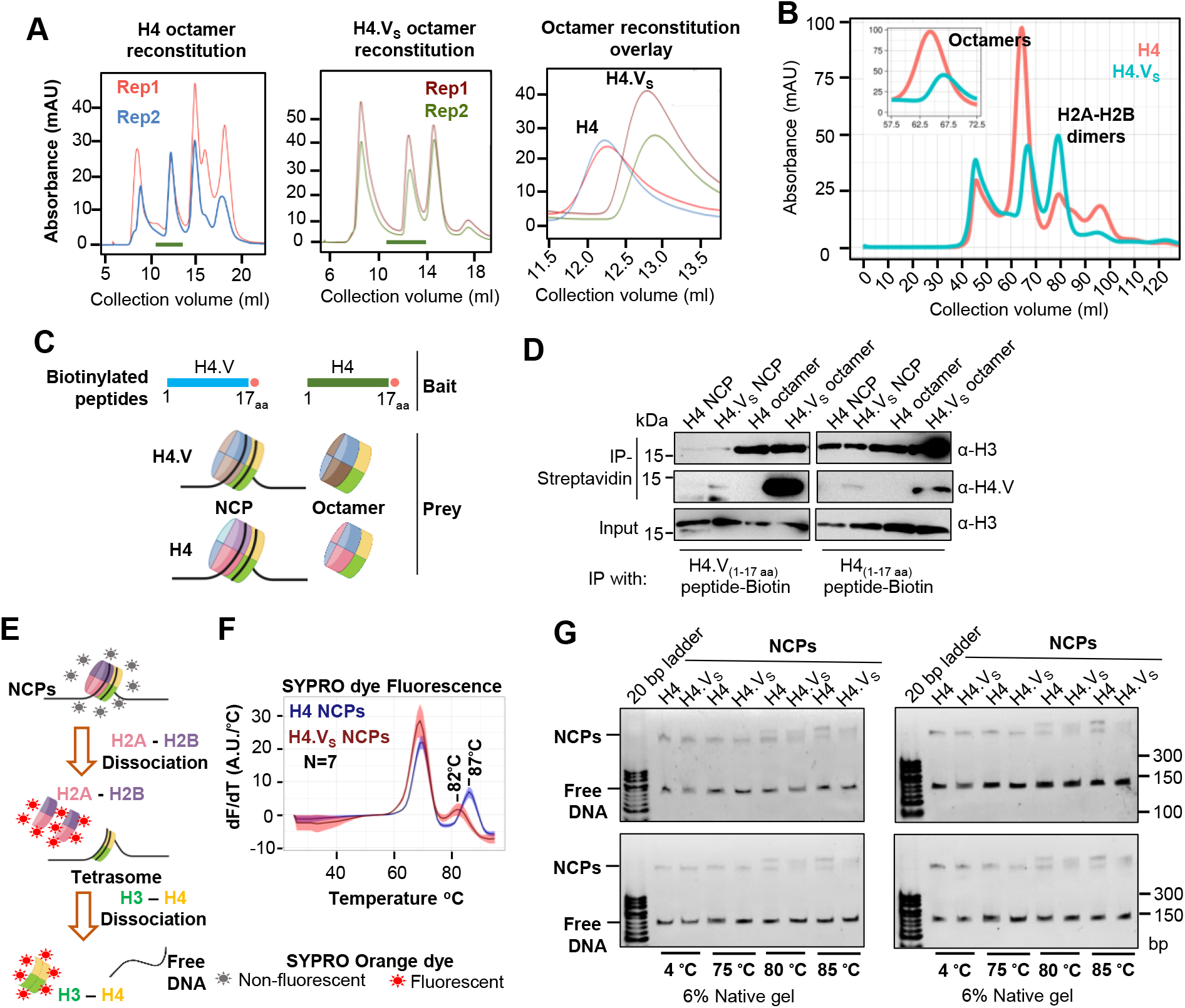
H4.V confers specific properties to nucleosomes. (A) Gel filtration chromatograms (24 ml S200 column) of the serially dialysed histone refolding mix containing H4 or H4.V_S_ to purify octamer fractions (green bars) from two independent experiments. Overlay of the chromatograms, zoomed at the octamer elution fraction (green bars) is shown in right. (B) Chromatogram of the same in a 120 ml S200 column. Inset plot shows range where octamers were eluted. (C) Schematic for the pulldown using biotinylated peptides of H4 and H4.V (bait) with the NCPs or octamers (prey). (D) Immunoblots showing pulldown products probed with α-H3 or α-H4.V. 10% of the prey complexes used as input control. (E) Schematic depicting the stepwise thermal decay of the NCPs. Fluorescent and non-fluorescent versions of the SYPRO Orange dye are shown in red and grey, respectively. (F) Thermal decay rates plotted as a function of temperature. Shaded region signifies 95% confidence interval from 7 replicates. First peak at ∼68 ᵒC depicts H2A-H2B dissociation and second peak depicts H3-H4 dissociation. (G) Gel-mobility shift assays of heat-treated NCPs. Four independent replicates are shown.

To check if the H4.V tail variations modulated interactions between the tail and DNA or octamer, we performed a peptide pulldown assay, where the first 17 residues (comprising maximum variations) of H4.V or H4 (with a biotin label, used as bait) were allowed to interact with recombinantly purified H4.V_S_ and H4 containing octamers and NCPs (prey) (Fig 3C). If the H4 variant tail is competent in interacting with the octamer core or the DNA itself, it must pull down the prey molecules. Using H3 histone as the unbiased readout for the pulled-down species, we observed that even though H4 tail efficiently interacted with all the prey species, H4.V tail interacted only with the octamers but not with the nucleosomes (Fig 3D). This indicated that H4.V tail residues were inefficient in surpassing the negatively charged DNA shielding to interact with the octamer core within, and pointed out a critical difference with H4 tail.

To test if the H4.V_S_ NCPs display varied degrees of stability, we traced a thermal decay profile using a fluorometric dye that quantitatively assessed the two-step denaturation of nucleosomes (Fig 3E). H3-H4 tetramer complex of H4.V_S_ NCPs denatured at a 5 ᵒC lower point when compared to H4 NCP tetramer (Fig 3F) unlike the H2A-H2B dimers. We also verified the lower denaturing point of H4.Vs NCPs using gel mobility shift assays that quantified fraction of intact NCPs after heat treatment (Fig 3G). These assays confirmed that the variations in the H4.V resulted in NCPs that are biochemically and structurally distinct in terms of their dimensions, intra- and inter-nucleosomal interactions and stability, all of which have the potential to translate to specific chromatin properties.

### Perturbation of H4.V resulted in growth and reproductive defects

To establish the importance of H4V during plant development we compared wild type with mutant rice plants devoid of H4V. Loss of function of H4.V (*h4.v KO*) showed growth defects in the early stages of development and the seedlings were stunted (Fig 4A and B). We used artificial microRNA (amiR) to obtain plants with reduced H4.V levels (*h4.v kd*) and overexpression lines (OE) of the H4.V, H4 and H4.V_S_ to explore the dosage dependent effects of H4.V (Appendix Fig S6). Reproductive development of *h4.v KO* plants were comparable to WT plants. On the contrary, OE of the H4.V led to reduction in seed filling, likely due to effects of spurious incorporation of the variant in the young reproductive stages (Fig 4C). OE-H4 did not show significant reduction in seed setting rate. Notably, OE of H4.V_S_ showed significant reduction in seed setting rate similar to OE of H4.V suggesting variable N-terminal tail is sufficient to bring changes in regulatory properties of H4.V. When the seeds matured, *h4.v KO* produced smaller seeds even though they did not show seed setting defects (Fig 4D and E). Transcriptome profiling in seedlings and leaves of the *h4.v KO*, OE-H4.V and OE-H4 showed large number of genes were mis-regulated in the H4.V perturbed lines (Fig 4F). Although the H4.V occupancy sites showed overlap with the H3K9me2-rich repressed constitutive heterochromatin (Fig 1K), a de-repression of transposons and repeats in these loci were not observed in the *h4.v KO* (Appendix Fig S7A), indicating H4.V might have specific roles in protein coding regions. In agreement with this, variations in neither the repeat derived small(s) RNAs nor DNA methylation status of LINE1 LTR-transposon that regulate the transcriptional silencing (Hari Sundar G *et al*, 2023) were observed in sRNA northern blots or methylation sensitive Southern blot (Appendix Figs S7B and C). It is possible that H4.V bound regions might show de-repression under specific conditions. These results suggest that although H4.V is predominantly located at heterochromatin it does not affect expression of transposons. However, it affects indirectly expression of protein coding genes and in the regulation of growth and reproductive development.

**Figure 4.**
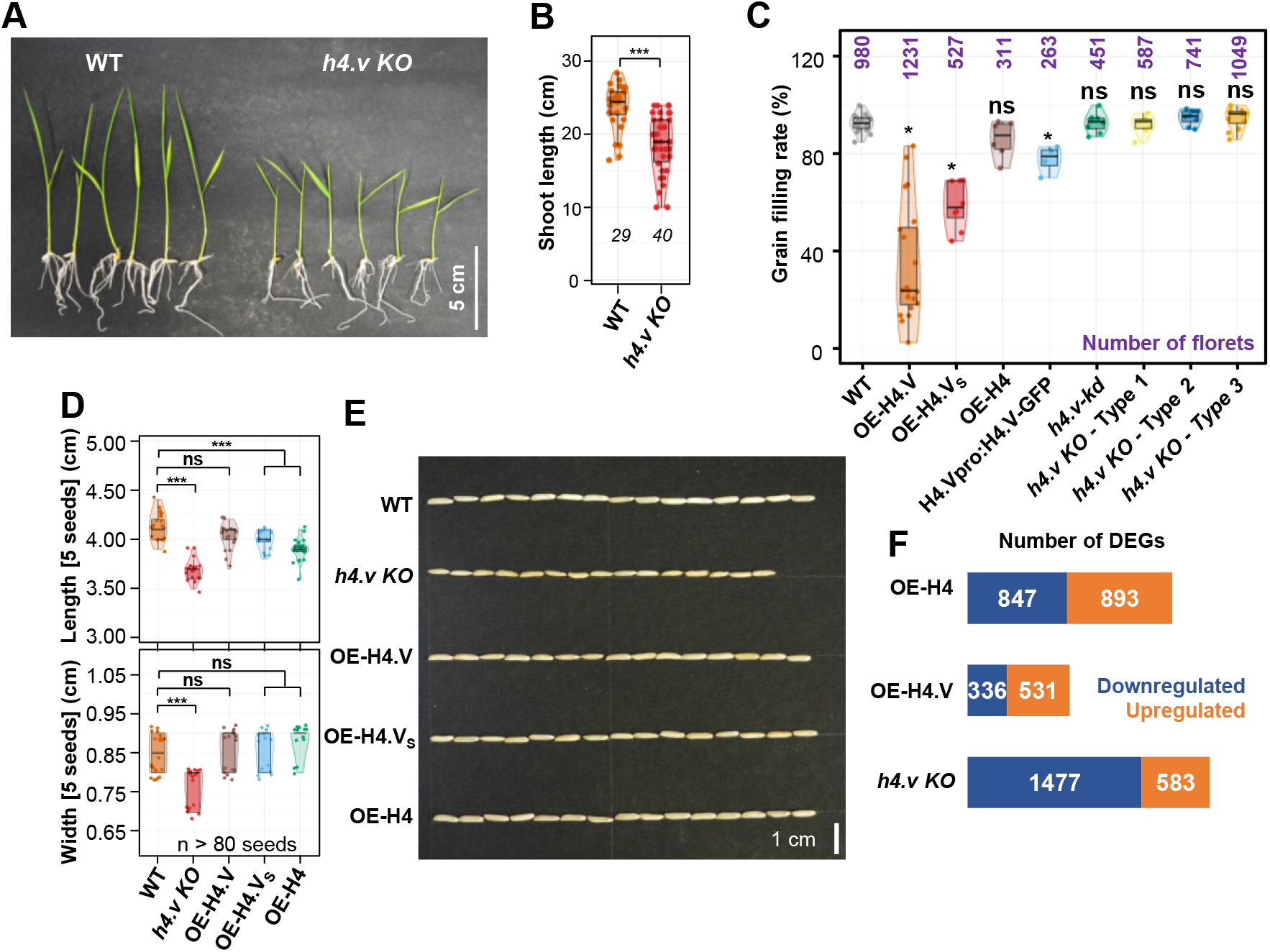
Histone H4.V perturbation leads to phenotypic defects largely attributable to gene mis-regulation. (A) Seedlings images of WT and *h4.v KO* plants. (B) Box-violin plots showing shoot length distribution of WT and *h4.v KO* plants. Number of plants taken is mentioned in italics (C) Box-violin plots showing percentage of filled grains. Dots represent data from individual panicles. (D) Box-violin plots showing seed size distribution taken 5 at a time. Dots represent dimensions of 5 seeds. At least 80 seeds were taken for analysis. (E) Representative image of 15 seeds from different genotypes. (F) Stacked bar plots showing number of DEGs in different genotypes. (B-D) Two-tailed Student’s *t*-test was used for statistical comparison. (*) p-value< 0.05, (ns) non-significant.

### H4.V prevented aberrant salt-stress like transcriptome

Histone variants were implicated in stress responses in plants (Nunez-Vazquez *et al*, 2022). Most of the stress responses in plants are mediated through regulation selected master transcription factors, including WRKYs, MYBs, NAC etc. (Javed *et al*, 2020). Differentially expressed genes (DEGs) in the *h4.v KO* were enriched with major TFs involved in stress responses (Fig 5A). Significantly downregulated genes were specifically enriched for the genes that respond to various stresses (Fig 5B). In order to identify the specific type of stress that elicits a transcriptional program that resembles *h4.v KO*, we analysed publicly available datasets (Kawahara *et al*, 2016) and performed a principal component analysis (PCA) for the DEGs identified in *h4.v KO* (Fig 5C). These analyses revealed that the *h4.v KO* DEGs respond most similarly to the salt stress response (Fig 5C and Appendix Fig S8A). The same response was also evident in the DEGs identified upon salt stress treatment (Fig 5D and Appendix Fig S8B).We performed salt stress on wild type and *h4.v KO* plants and observed that 65% of the DEGs in *h4.v KO* were commonly mis-regulated upon salt stress (Fig 5E). Further, we compared the gene expression profiles of *h4.v KO* DEGs in WT and *h4.v KO* plants with and without salt stress. We observed that *h4.v KO* DEGs showed similar status in wild type plants under salt stress (Fig 5F). Surprisingly, salt stressed *h4.v KO* plants exhibited similar trend of gene expression as that of the stressed WT plants (Fig 5F), suggesting absence of H4.V triggered a precocious salt-stress like transcriptional profile. Candidate genes that are well-established master-controllers of salt stress responses in rice (Liu *et al*, 2022; Horie *et al*, 2012) via various signaling modes have also been perturbed in *h4.v KO* similar to salt stress (Appendix Fig S9). To quantitatively map the correlation between salt stressed state and *h4.v KO*, we plotted scatter plots of degrees of variation of the expression of DEGs across the two states. We observed a statistically significant positive correlation effect (Fig 5G and Appendix Fig S8C supporting the notion that H4.V is indeed involved in preventing salt stress like transcriptional state. As *h4.v KO* plants exhibited salt stress-like transcriptome, we subjected the H4.V perturbed lines to prolonged salt stress (Methods). Wild type rice plants, under 120 mM salt stress show extended primary roots but reduced secondary roots and stunted shoots and this behaviour is a proxy for the responses to the salt stress (Pachamuthu *et al*, 2022). Our assays showed that *h4v KO* plants poorly responded to salt stress (Fig 5H) in addition to being stunted in growth. In summary, our observations suggested that H4.V is necessary for controlling transcription of necessary salt-responsive TFs and genes aiding in coping with the stress.

**Figure 5.**
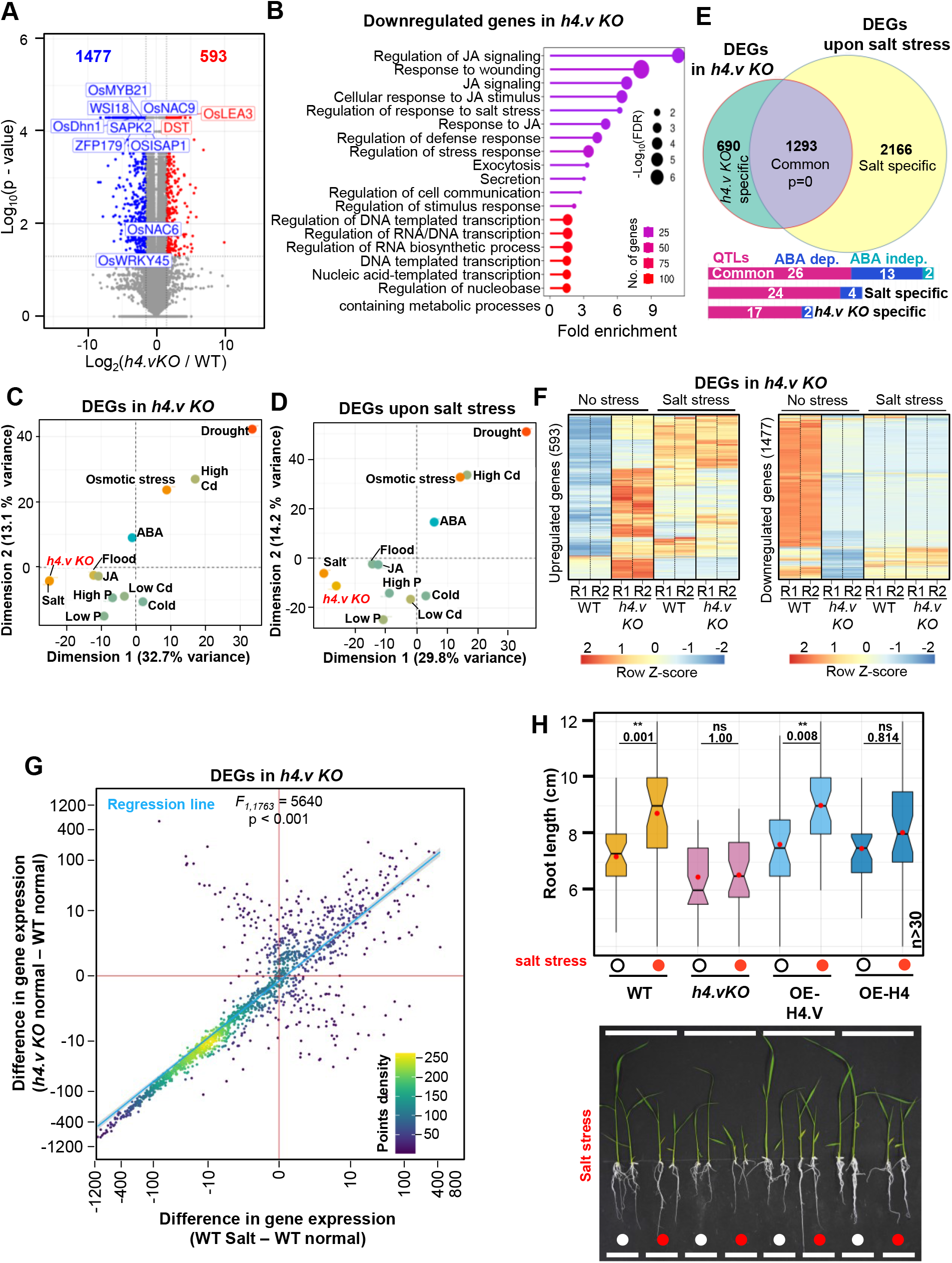
KO of H4.V exhibits a transcriptome similar to salt stressed state. (A) Volcano plot showing the DEGs in *h4.vKO* seedlings. Blue dots represent downregulated and red dots represent upregulated genes. Labelled genes are stress-responsive. (B) GO enrichment categories among the downregulated genes. FDR-false discovery rate. (C-D) PCA plots of selected stress dependent transcriptomes (full list in Appendix Fig S8) for the *h4.v KO* DEGs (C) and salt stress DEGs (D). The first two principal dimensions are plotted and the percentage of variance explained is indicated. (E) Venn diagram representing the overlap between *h4.v KO* and salt stress DEGs. Hypergeometric test was used for statistical testing of overlaps. Bar plots show no. of genes represented in each category with salt-linked QTLs, ABA dependent/independent salt responsive genes. (F) Clustered heatmaps showing variation in expression across WT and *h4.v KO*, with and without salt stress across *h4.v KO* DEGs. (G) Density scatter plots showing resemblance of *h4.v KO* with salt stress transcriptomes over *h4.v KO* DEGs. Points density is color coded. Blue line represents the linear regression fit line, and grey shade represents the 95% confidence interval. Difference in gene expression is plotted with x- and y-axes are scaled to inverse sine hyperbolic function. F-test was used for statistical testing of the dependence of two perturbations (salt stress and h4.vKO). (H) Box plots showing root length distribution without and with salt stress (red circles) across genotypes. Red dots within the boxes represent mean of the values. At least 30 plants were phenotyped for each condition/genotype. Student’s *t*-test was used for statistical testing. P-values are mentioned across comparisons. Representative image of the plants used for phenotyping are shown below.

### Salt stress induced increased H4.V occupancy

In order to check the contribution of H4.V upon salt stress, we profiled the H4.V abundance and occupancy upon salt stress. We observed increased fluorescence signal form the H4.V-GFP translational fusion lines upon salt stress suggesting increased accumulation of the H4.V (Fig 6A). Furthermore, spatial patterning of H4.V in the IFL micrographs revealed wider punctate occupancy in the salt stressed nuclei when compared with the unstressed state (Fig 6B). To check if the H4.V occupancy was altered upon salt stress, we analysed H4.V profiles with and without salt stress. This analysis revealed newer occupancy sites corroborating the microscopic observations (Fig 6C). We also performed differential enrichment analyses of H4.V binding sites upon salt stress (Methods) which identified the expanded-occupancy of the H4.V upon salt stress (Figs 6D and E). These results agree with the transcriptional mis-regulation profiles (Fig 5), and suggest that H4.V responds to the salt stress and its relative occupancy might modulate the gene expression pattern promoting salt dependent responses.

**Figure 6.**
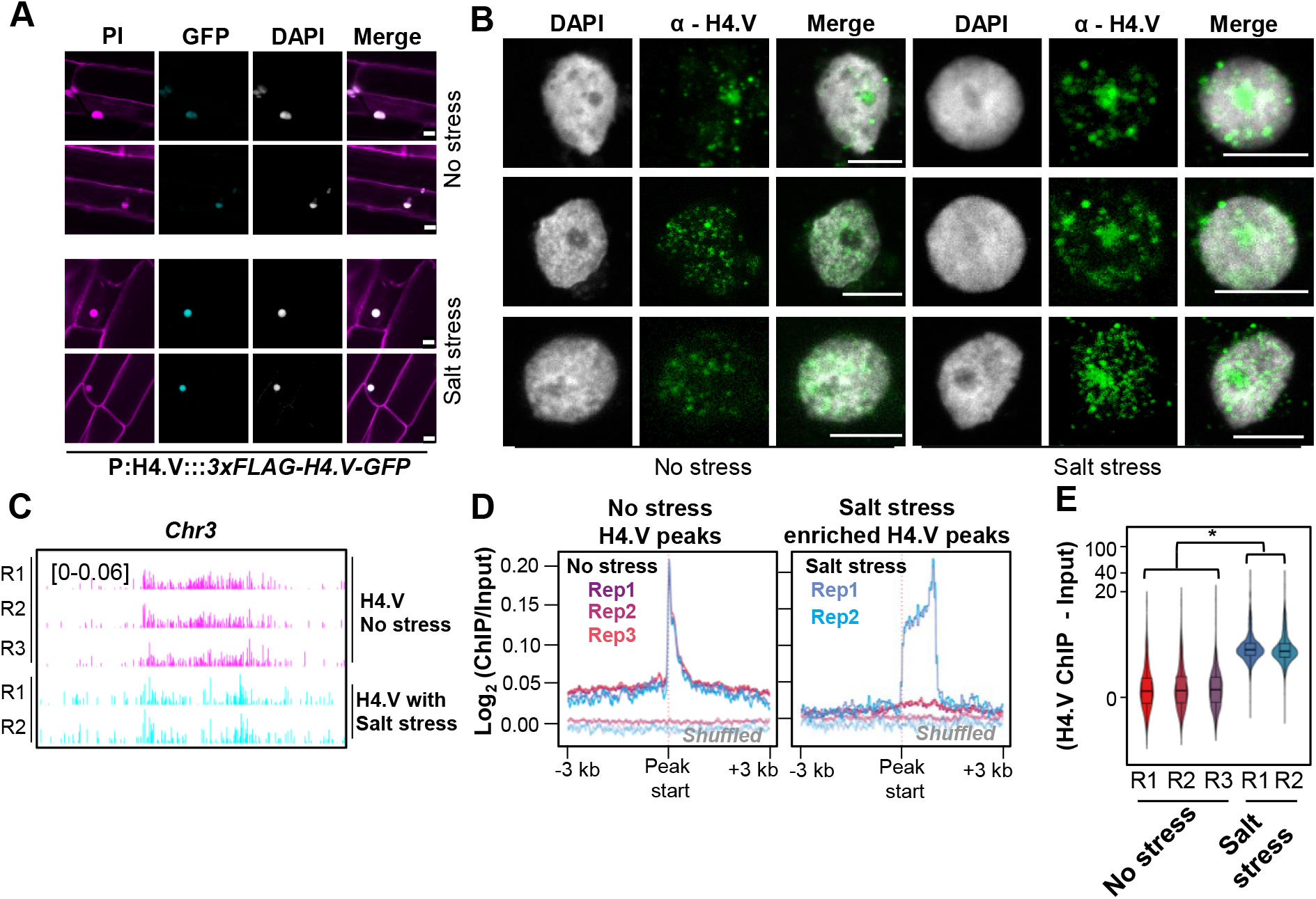
H4.V occupancy redistributes upon salt stress. (A) Fluorescent micrographs of seedling roots expressing 3xFLAG-H4.V-GFP. Seedlings grown without and with salt stress are shown. (B) IFL images of nuclei stained using α-H4.V antibody. Nuclei isolated from seedlings grown without and with salt stress are shown. (C) Chromosome-wide genome browser screenshots showing ChIP enrichment (square brackets) of H4.V without and with salt stress. (D) Metaplots showing enrichment of H4.V with and without salt stress. (E) Box-violin plots showing enrichment of H4.V over the salt-stress enriched peaks (n=1700). Y-axis is scaled to inverse sine hyperbolic function. Wilcoxon signed rank sum test was used for testing statistical significance. (*) p-value < 0.001. (A-B) Details as in Fig 1E.

### H4.V affects H4K5Ac deposition in response to salt stress

Chromatin readouts, in terms of transcription or accessibility, is a combined effect of several layers of epigenetic modules and interaction partners. H4.V confers unique nucleosome properties (Figs 2 and 3) that might facilitate or hinder other epigenetic layers by acting as guides of gene regulation. To test the link between H4.V and other chromatin modifications, we obtained several histone modification ChIP-seq datasets from rice seedlings and called for enriched peaks (Methods, Appendix Dataset S1). In addition to H3K9me2, H4K5Ac marks also showed significant overlap with the H4.V peaks that are predominantly different from the H3K9me2 overlaps (Fig 7A). Other known gene regulatory histone marks did not enrich at the H4.V regions. It is important to note that H4.V itself is incapable of undergoing H4K5Ac modification as it lacks the 5^th^ Lysine residue (Fig 1A). This implied that H4.V might be forming heterotypic nucleosomes with canonical H4 that can undergo H4K5Ac and H4.V modulated the efficiency of H4K5Ac modification. To check the influence of H4.V on the H4K5Ac marks, we performed ChIP-seq to detect H4K5Ac in the WT and *h4.v KO* seedlings with and without salt stress. H4K5Ac is a known pioneer regulator of salt stress response in plants and it is under the control of several stress-responsive histone deacetylases and acetyl transferases (Ueda *et al*, 2017; Ullah *et al*, 2020; Zheng *et al*, 2019; Feng *et al*, 2022; Zheng *et al*, 2021). We found that in *h4.v KO,* H4K5Ac marks were absent at the sites originally occupied by H4.V and H4K5Ac (Figs 7B and C). Surprisingly, upon salt stress, the same sites lost H4K5Ac as seen in *h4.v KO* (Fig 7B). This remarkable similarity of H4K5Ac dynamics in *h4.v KO* and upon salt stress pre-empted the notion that H4.V is necessary for the H4K5Ac modulation upon salt stress. *h4.v KO* did not cause global loss of the H4K5Ac marks and bulk of the marks were retained in the *h4.v KO* (Fig 7D and Appendix Fig S10). To understand the genome-wide modulation of the H4K5Ac marks due to H4.V, we overlapped the H4K5Ac peaks in WT and *h4.v KO*. We found 35% of the H4K5Ac peaks in WT were lost in *h4.v KO* (H4K5Ac peaks lost in *h4.v KO*, 15464 peaks) and 9689 new peaks emerged in *h4.v KO* (Figs 7E and F), definitively suggesting the indirect effect of H4.V on the distribution of H4K5Ac marks. Surprisingly, the H4K5Ac peaks that were lost in *h4.v KO* (15464 peaks) were also lost upon salt stress in WT plants and the newly emerged peaks in *h4.v KO* were the sites that showed enrichment upon salt stress (Figs 7E and F). To further validate similarity of H4K5Ac profiles between salt stress and *h4.v KO,* we overlapped the differentially enriched H4K5Ac peaks upon salt stress with the mis-localised H4K5Ac peaks in H4.V. Even though salt stress specific peaks outnumbered and did not colocalize with the *h4.v KO* specific H4K5Ac peaks, 71% (10974) of the salt stress eliminated H4K5Ac peaks colocalized with the peaks lost in *h4.v KO* (Fig 7E and F). On the other hand, H4K5Ac peaks that were retained in *h4.v KO* overlapped 93% of the salt-persistent H4K5Ac peaks and showed no change upon salt stress. This data suggested that the localisation of H4K5Ac was dependent on the H4.V status and the H4.V perturbation sufficiently mimicked the H4K5Ac profile generated upon salt stress. Since we observed a salt-stress like transcriptome in *h4.v KO* (Fig 5F), we speculated that the gene mis-expression was related to changes in the profiles of H4K5Ac. To test this, we identified the genes overlapping with the H4K5Ac peaks that were lost in *h4.v KO* (4793 genes) and found they were mis-regulated upon *h4.v KO* (Fig 7G). Further, of the 4793 genes, we picked the genes overlapping with H4K5Ac peaks that were also lost upon salt stress (2071 genes), which also showed similar but exacerbated response (Fig 7G). The extent of mis-regulation of genes upon salt and *h4.v KO* for these genes were also linearly dependent (Fig 7H), further validating our model of gene mis-expression occurring via H4K5Ac marks. Taken together, these results suggested a previously uncharacterized regulator of H4K5Ac marks, H4.V, whose occupancy dictated the H4K5Ac marks and hence the gene expression necessary for salt stress responses. The above results demonstrated that the genes identified as salt stress mediators including well known TFs such as WRKY45, NACs and MYBs (Appendix Fig S9), regulation of which was previously ascribed to histone PTMs such as H4K5Ac (Ueda *et al*, 2017; Ueda & Seki, 2020), are also under an additional layer of control by the newly identified H4.V.

**Figure 7.**
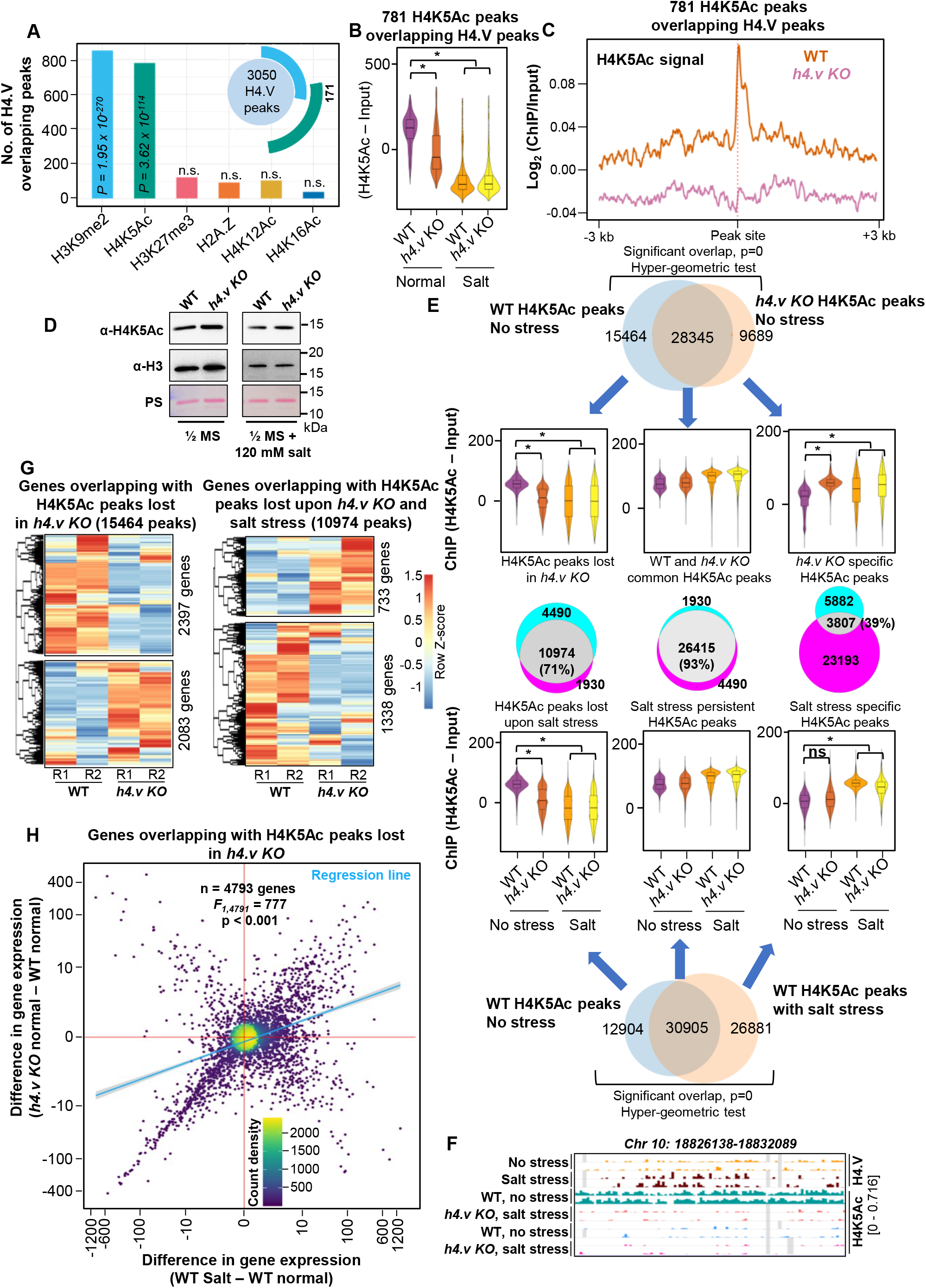
H4 variant occupancy determines occupancy of H4K5Ac marks thereby regulating genes. (A) Bar plots showing number of overlaps of H4.V peaks with peak sets of other histone marks. Significance of overlap was tested using hyper-geometric test and p-values above 0.05 are mentioned. The inset plot shows number of peaks that are overlapping with H4.V and other histone marks (overlaps more than 20 are shown). (B) Box-violin plots depicting enrichment of H4K5Ac marks over the H4K5Ac peaks that overlap with the H4.V marks. (C) Metaplot showing enrichment of the H4K5Ac mark distribution over the H4K5Ac peaks that overlap with the H4.V marks. (D) Immunoblots showing the global levels of H4K5Ac marks without and with salt stress. H3 levels and Ponceau stained blots served as loading controls. (E) Venn diagrams and H4K5Ac enrichment profiles for the overlaps and cross-overlaps of H4K5Ac peaks without and with stress and in *h4.v KO*. Significance of overlaps were tested using hyper-geometric test and that of the enrichment differences were computed using Wilcoxon signed rank sum test. Percentage of overlap is shown. (F) Genome browser screenshot showing ChIP enrichment (square brackets) of H4.V and H4K5Ac marks. (G) Heatmaps showing the expression profiles of genes overlapping with perturbed H4K5Ac marks. Number of genes in each hierarchical cluster is mentioned. (H) Density scatter plots showing resemblance of gene expression profiles upon *h4.v KO* and salt stress for the genes that overlap with H4K5Ac perturbation. Additional details are in Fig 4G. (B and E) Wilcoxon signed rank sum test was used for testing statistical significance. (*) p-value <0.001.

## Discussion

Histones variants are consequence of evolutionary events that resulted in novel modifications that acquired new properties. There have been several instances of independent evolution of histone variants with similar modifications in H2A and H3 in plants and animals implying the importance and utility of these modifications in conserved cellular roles (Talbert & Henikoff, 2017; Yelagandula *et al*, 2014; Malik & Henikoff, 2003).

Histone variants can incorporate residue substitutions and motif additions to the canonical histones and innovate at least 4 specific downstream functions (Osakabe *et al*, 2018; Yelagandula *et al*, 2014; Schmücker *et al*, 2021). These include a) abrogation/facilitation of PTMs, b) drive assembly of homotypic/heterotypic nucleosomes arising out of variable amino acid sequences, c) modification of the physico-chemical and structural properties of the nucleosomes and d) modulation of the spatio-temporal interactions with distant nucleosomes or chromatin regulators. Among all the histones, H4 is the slowest evolving histone as it contacts with several residues in the histone octamer complex and the H4 tail is a hotspot of several regulatory PTMs keeping it under strong negative selection (Malik & Henikoff, 2003). H4.V has most of the modifications in the N-terminal tail that affects PTMs but conserves the residues in the core of the histone enabling octamer’s core interactions. H4.V hence acts as an effective switch to globally change the histone PTMs. Among the H4.V specific modifications, PTM sites K5, K8, K14, K16 and K20 are abrogated and these are linked predominantly to abiotic stress responses (Nunez-Vazquez *et al*, 2022; Pavangadkar *et al*, 2010; Roca Paixão *et al*, 2019). These modified N-terminal tail residues, if acetylated, directly modulates accessibility to DNA, thus promoting a combined histone-TF code (Wakamori *et al*, 2015; Michael *et al*, 2023).

Adding to the regulatory potential, the H4-tail residues have unique properties that contribute to the modulation of the spatio-temporal and structural interactions with other proximal or distal nucleosomes and DNA (Kan *et al*, 2009; Stormberg *et al*, 2021). A closer look at the amino acid compositions of human and rice histones showed amino acid variations in C-terminal region of rice H2A, which likely contributes to its weakened interactions with H3 αN helix as observed in human NCP (Luger *et al*, 1997). The surface charge properties of the core histone regions in the nucleosomes (like the H2A-H2B acidic patch) often mediates interactions with chromatin remodellers and histone readers/modifiers. A comparison of the electrostatic surface features of rice and human nucleosomes indicated that the canonical rice nucleosome has an acidic patch that is spread beyond H2A-H2B and extends to H3/H4 regions (Fig 2G). Interestingly, the acidic patch distribution of the H4.Vs nucleosome is noticeably different as compared to the canonical nucleosome. The variation mainly appears to be due to the Glu to Lys substitution on H4.V at position 53 (Fig 2G). Overall, our structural characterisations suggest that the H4.Vs NCP’s protection of entry/exit DNA from unwrapping and the distinct acidic patch features might contribute to modulating key interactions with chromatin remodellers and chromatin readers/writers involved in specific reprogramming of transcriptional regulation (Fig 2). It is interesting to note that a recent study uncovered that a chromatin remodeller, Decrease in DNA Methylation1 (DDM1), binds to the H4 tail residues anchoring *via* non-acetylated residues (Lee *et al*, 2023; Osakabe *et al*, 2023). Since H4.V lacks the residues that can be modified, it would be interesting to know if DDM1 can interact with H4.V NCPs and engage in epigenetic modifications. We observed atypical properties of H4.V containing nucleosomes (Fig 3) in terms of the stability, nucleosome-DNA interactions, condensed octamer core and accessibility, indicating H4.V might be contributing to major classes of alterations including interactions with remodellers and modifiers.

H4.V is found only in the *Oryza* genera among the sequenced species. Several histone variants are also restricted to specific clades as they are continuously evolving and are recent innovations necessitated by the unique environmental niches where these organisms evolve. For instance, *Hominidae* family restricted H4G and divergence of histones among kinetoplastids and Mycetazoans confer unique functional roles commensurate with the genomic divergence (Long *et al*, 2019; Alsford & Horn, 2004; Poulet *et al*, 2021). Early-reproductive tissue specific exclusion of the H4.V is functionally significant as ectopic OE of the variant in reproductive tissues caused reproductive defects (Fig 4). Similarly, in *Arabidopsis* seed and sperm cell specific H2B.S, when ectopically expressed in vegetative tissues, caused aberrant chromatin compaction (Buttress *et al*, 2022). In addition, *h4.v KO* showed smaller seeds and stunted growth. Possible roles of H4.V in tissue specific chromatin regulation is an avenue for future investigation. Wild relatives of *Oryza* members propagate predominantly by vegetative means by having perennial rhizomes (Tong *et al*, 2023) and it is possible that exclusion of H4.V in sexual reproductive tissues is an indication of its specific role in vegetative tissues and in regulating chromatin under stresses.

About 20% of the arable land is already affected by salinity (Butcher *et al*, 2016). Understanding the salt tolerance mechanism of crop plants that have strains or varieties with this ability in comparison with susceptible lines provides greater confidence to find a way to apply the knowledge to breed better crops. Most monocots including *Oryza* members have a mechanism mostly to exclude salt from their semi-aquatic environments. Key steps in salt tolerance mechanism include a) reduced uptake - apoplastic barriers such as casparian bands and suberin lamellae in roots b) exclusion - transporting Na+ out of the root or into the vacuoles by Na+/H+/K+ transporters; or c) tissue tolerance – osmotic adjustments by accumulation of ions, solutes and osmolytes (Reddy *et al*, 2017; Horie *et al*, 2012; van Zelm *et al*, 2020). These steps in salt tolerance involve signaling both dependent and independent on abscisic acid (ABA) (Kumar *et al*, 2013). The stunted growth phenotypes of the *h4.v KO* are likely due to the aberrant ABA signaling involved in salt like responses. The responses to salt stress are spatially and temporally distinct with shoot and root dynamically regulating transport (early) followed by metabolic biosynthesis and transport and finally structural and morphological changes. This regulated multi-pronged response requires interplay among several TFs belonging to WRKY, MYB, bZIP and NAC subtypes. These TFs can bind to hundreds of gene promoters synergistically and brings about the salt responses. In *h4.v KO*, we documented mis-regulation of NAC, MYB and WRKY type TFs that showed similar responses like that of plants under salt stress (Supplementary Fig S9). Enrichment of H4K5Ac marks at these genes were also reduced, suggesting H4K5Ac mediated transcriptional mis-regulation poised by H4.V marking of these genes. These results support the idea of evolution of H4.V as an upstream regulator of these salt exclusion mechanisms that need to be employed seasonally in a semi-aquatic species such as *Oryza* members. Such a mechanism operating at the chromatin level might have allowed successful adaptation of *Oryza* genera members to diverse water limiting conditions encountered routinely, both in their natural habitats or in the agricultural set up in which they were domesticated.

## Materials and methods

### Plant material

Unless specified otherwise, *indica* rice variety (*O. sativa indica sp.*) Pusa Basmati 1 (PB1) was used for analyses. Plants were grown in a greenhouse maintained at 28ᵒC with a natural day-night cycle. Other rice varieties – *O. nivara, O. rufipogon* and *O. sativa indica* variety pokkali were also grown in the same conditions.

### Cloning, plasmid construction and transgenic plant generation

For generation of CRISPR Cas9 mediated knockouts, gRNA (5’-gucuuucgcggcuacaucca-3’) coding fragment was cloned into pRGEB32 using HindIII and BsaI sites (Xie & Yang, 2013). Rice H4.V and H4 CDS were amplified (primers in Appendix Table S4) from cDNA prepared from seedling RNA with Superscript III reverse transcriptase (Invitrogen). H4.V_S_ CDS was prepared by Gibson ligation of the PCR fragments from H4.V and H4. For the amiR-mediated *h4.v kd*, amiRNA (5’-uagacaauccgaucgugccua-3’) was encoded by swapping the *Osa*miR528 coding region in pNW55 using WMD3 tool (Ossowski *et al*, 2008). For OE lines and the *h4.v kd* lines, CDS or the precursor of amiR region were amplified and cloned into a derivative of pCAMBIA1300 plasmid between maize Ubiquitin promoter and 35S-terminator sequences. All the binary plasmids were mobilised into *Agrobacterium tumefaciens* strain LBA4404 harbouring extra-virulence plasmid pSB1. Embryogenic rice calli were infected with *Agrobacterium* strain containing binary plasmids using established methods (Hiei *et al*, 1994; Sridevi *et al*, 2003) and the calli were selected and regenerated on hygromycin containing medium.

### Grid preparation and cryo-EM data collection

Samples were prepared as previously described in (Migl *et al*, 2020). Prior to sample vitrification, samples were concentrated up to a DNA concentration of 900 ng/µl in 10 mM Tris pH 8.0, 50 mM NaCl, 1 mM EDTA. H4 NCPs were crosslinked with 0.005 % (V/V) glutaraldehyde for 5 min. The crosslinking reaction was quenched by adding 50 mM Tris pH 8.0, allowing it to react for 1 h. Right before grid preparation, Tween-20 was added to a final concentration of 0.005 %. H4 NCPs were applied to Quantifoil R2/1 200-mesh grids coated with 2 nm of carbon (Quantifoil). H4.V_S_ were applied to unsupported Quantifoil R2/1 200-mesh grids (Quantifoil). Grids were glow-discharged for 20 s in the presence of air using a current of 20 mA. 4.5 μl of sample were used per grid. Grids were blotted for 2.5 s at 10 °C and 95 % humidity using a Leica EM GP plunger (Leica). Data for NCPs was acquired at the Cryo-EM facility of the Gene Centre in Munich. Datasets were acquired on a Titan Krios transmission electron microscope operating at 300 keV and equipped with a Falcon4i direct electron detector and a Selectris X Energy Filter (energy slit width of 5 eV) (ThermoFisher). The parameters used for data collection are provided in Appendix Table S6. Automated data collection was done using the Smart EPU software (ThermoFisher). MotionCor2 was used to perform motion correction of the movies (Zheng *et al*, 2017). Motion corrected micrographs were processed using CryoSPARC v4.0.2 (Punjani *et al*, 2017). For both datasets, manual picking and further 2D classification were used to obtain 2D classes presenting defined secondary structure features. These particles were used as a template for template picking. After several rounds of 2D classification, resulting particles were used to train Topaz, which was subsequently utilised to enrich the particle dataset. Particles were extracted using a 356-px box size of (down sampled to 80-px box size). Further rounds of 2D classification were performed until 2D classes presented high resolution features. This pipeline yielded 177,136 particles for the H4 NCP dataset and 161,580 particles for the H4.V_S_ NCP dataset. Per dataset, several *ab initio* models were generated and subjected to heterogeneous refinement. Particles corresponding to the best model were re-extracted using a box size of 300 px for the H4 NCP dataset and 356 px for the H4.V_S_ NCP dataset. For the Canonical NCP dataset, two *ab initio* models were further generated and subjected to heterogeneous refinement. Particles for the best model were cleaned using 2D classification and a final model was generated using 140,877 particles by *ab initio* model generation followed by non-uniform refinement (Punjani *et al*, 2020). This yielded a final 3D reconstruction at 3.62 Å resolution (Fourier shell correlation (FSC) = 0.143) for the H4 NCP dataset. For the H4.V_S_ dataset, one *ab initio* model was generated after particle re-extraction (148,277 particles). This model was refined using non-uniform refinement, which yielded a final 3D reconstruction at 3.64 Å resolution (FSC = 0.143). For both NCP datasets, the 3D Flex pipeline in CryoSPARC (Punjani & Fleet, 2023), was further used as a refinement tool to resolve flexible regions. This resulted in two final maps at 3.6 Å resolution (FSC = 0.143) for both the NCPs. Initial models comprising rice histone octamers were obtained using Alphafold2. NCPs were assembled combining AlphaFold2-generated rice histone octamers and Widom 601 DNA structure from human NCP (PDB: 7XD1). The models were placed in the density map obtained from the 3D Flex pipeline using the rigid fitting option available in ChimeraX. Models were refined using real-space refinement in the Phenix suite of programs. Model building was performed using Coot.

### Multiple sequence alignment and phylogeny

Histone H4 sequences (Appendix Table S1) were aligned using ClustalW and the phylogeny was visualised using iTOL (Letunic & Bork, 2021).

### Reverse transcriptase – quantitative polymerase chain reaction (RT-qPCR)

TRIzol reagent (Invitrogen) was used to extract total RNA from plant tissues and around 1.5 µg of DNAseA treated RNA was taken for cDNA synthesis using oligo-dT and random hexamers mix as primers (Invitrogen - SuperScript III RT kit). The cDNA template was used for qPCRs using SYBR green (Solis Biodyne – 5x HOT Firepol Evagreen qPCR master mix) with *GAPDH* (LOC_Os04g40950) or *Actin* (LOC_Os03g50885) as an internal control.

### sRNA northern hybridisation

sRNA northern hybridisation was performed as described earlier (Shivaprasad *et al*, 2012; Tirumalai *et al*, 2020). Around 8 µg of total RNA was electrophoresed on a denaturing gel, blotted onto Hybond N+ membrane (Cytiva) and UV crosslinked. The membrane was hybridised with radioactively labelled probes (T4 PNK, M0201, New England Biolabs (NEB)) in Ultrahyb buffer at 35 ᵒC and exposed to a phosphor screen that was scanned using Typhoon scanner (Cytiva).

### Southern hybridisation

Southern hybridisation was performed as described earlier (Hari Sundar G & Shivaprasad, 2022; Ramanathan & Veluthambi, 1995). Probes were PCR amplified and internally labelled using Rediprime labelling kit radioactively (Cytiva).

### Antibody generation

Polyclonal antibodies against H4.V were raised against a peptide CAPRSVAISGRGTSGA in rabbit and antigen-affinity purified by Lifetein LLC, USA.

### IFL and microscopy

IFL microscopy was performed as described earlier (Yelagandula *et al*, 2014; Hari Sundar G *et al*, 2023). Crosslinked seedlings were chopped and isolated nuclei were crosslinked on a positively charged slide. The nuclei are stained using α-H4.V (1:100), α-H4K5Ac (Merck 07-327, 1:200) and α-H3K9me2 (Abcam ab1220, 1:100) antibodies and detected using fluorescently tagged secondary antibodies (Invitrogen, goat raised anti-mouse 488 and Anti-rabbit 555). Nuclei were counterstained with DAPI and imaged on a confocal microscope (Olympus FV3000).

### ChIP-seq and sequence analysis

ChIP was performed from 1.5 g crosslinked rice seedlings as described earlier (Saleh *et al*, 2008; Hari Sundar G *et al*, 2023). Sheared chromatin was incubated with 10 µg of α-H4.V or 5 µg of α-H4K5Ac (Merck 07-327) or 4 µg of α-H4 (Abcam ab10158) overnight at 4 ᵒC and further incubated with protein-G Dynabeads (Thermo Fisher Scientific) for four hours. The purified IP products were prepared into a library using NEBNext Ultra II DNA library prep kit (NEB E7103) using manufacturer’s protocol. The libraries were sequenced on an Illumina HiSeq 2500 or NovaSeq 6000 (Appendix Table S2).

Cutadapt trimmed reads were mapped to IRGSP1.0 genome using Bowtie 2 (Langmead & Salzberg, 2012; Martin, 2011) with the following parameters: -v 1 -k 1 -y -a -best - strata. Alignments depicting PCR duplicates were removed and the alignments were converted to enrichment (log_2_) coverage tracks normalising to sheared input DNA using deepTools (Ramírez *et al*, 2014). Coverage signals over regions of interest were computed using ComputeMatrix (deepTools) and plotted using custom ggplot2 scripts in R (Wickham, 2011). ChIP peaks were called using MACS2 (Zhang *et al*, 2008) with broad peak calling with respect to the input DNA control and significant peaks with 1.5 fold enrichment were considered as true peaks. Peaks obtained were analysed and merged across replicates using BEDTools (Quinlan & Hall, 2010). ChIPseekeR (Yu *et al*, 2015) was used to annotate the peaks, overlap Venn diagrams were generated using Intervene (Khan & Mathelier, 2017). ChIP-seq datasets that are publicly available (Appendix Table S3) from previous studies were analysed in the same way and the peaks called from ChIP datasets are in Appendix Dataset S1. Differential enrichment of histone peaks across genotypes and conditions were performed using Diffreps (Shen *et al*, 2013) with the parameters described earlier (Hari Sundar G *et al*, 2023) (Appendix Dataset S1).

### Recombinant histone purification and nucleosome reconstitution

Rice histones purification, octamer reconstitution and nucleosome titration were done as described previously with following modifications (Abad *et al*, 2019; Luger *et al*, 1999). Rice histones CDS were codon optimized and synthesised (Thermo Fisher Scientific). The histone sequences, inducible expression plasmids, *E. coli* expression strain, expression conditions for each histone and Widom 601 NPS (147 bp and 188 bp) are provided in Appendix Table S5. The NCPs’ quality was estimated by electrophoresing on a native 6% acrylamide gel in 0.5x TBE buffer.

### NCPs MNase digestion, sequencing and analyses

For MNase digestion, 7 µg (DNA mass) of NCPs (with 188bp NPS) or free DNA was incubated with 5 Kunitz units of MNase (NEB M0247) in a reaction buffer (30 mM Tris pH 8.0, 5 mM CaCl2, 1.5 mM DTT, 0.1 mg/ml BSA) at 37 ᵒC in a reaction volume of 60 µl. Reaction was allowed to proceed for the mentioned time points and from the reaction mix 15 µl was withdrawn periodically and mixed with equal volume of 2x deproteinization buffer (20 mM Tris pH 8.0, 80 mM EDTA, 80 mM EGTA, 0.25% SDS, 0.5 mg/ml proteinase K) to stop the reaction. About 5 µl from this sample was electrophoresed on a 10% Acrylamide gel at 100V for 2 hours and stained with EtBr for visualisation. The remaining digestion products were purified and libraries were prepared like ChIP-seq samples and deep sequenced on Illumina NovaSeq 6000 in a 2×100 bp format. The obtained reads were adapter trimmed (Cutadapt), the overlapping paired end reads were stitched using FLASH (Magoč & Salzberg, 2011) and aligned to 188 bp NPS using Bowtie 2. Subsequent analyses and coverage plots were performed like ChIP-seq.

### Histone peptide pull-down assay

MyOne Streptavidin beads (40 µl per reaction) (Invitrogen, 65001) and 25 µg of biotinylated peptides (1-17 amino acids) were allowed to bind in a binding buffer (10 mM Tris pH8.0, 50 mM NaCl and 5 mM MgCl_2_) and subsequently washed in the same buffer. Peptide-bound beads were allowed to interact with 60 pmoles of NCPs (DNA estimation) in 0.5 ml binding buffer for 6 hr at 4 ᵒC. For the octamers, 250 pmoles of complex was allowed to interact in binding buffer with 2 M NaCl. Post-binding reaction, the beads were washed 4 times with corresponding binding buffers and eluted at 99 ᵒC in 50 µl Laemmli loading buffer. 35 µl of the precipitated products were immunoblotted.

### RNA extraction and transcriptome profiling

Total RNA was extracted using TRIzol from 14 d old seedlings and poly(A) enriched before library preparation (NEB E7490 and E7765). Libraries were sequenced on Illumina Hiseq 2500 in a 2×100 bp format. The obtained reads were adapter trimmed using Trimmomatic (Bolger *et al*, 2014) and mapped to IRGSP1.0 genome using HISAT2 (Kim *et al*, 2015). Cufflinks (Trapnell *et al*, 2012) was used to perform transcript abundance estimation and differential gene expression analyses. The DEGs were called with a p-value cut-off of 0.05 and absolute log_2_ (fold change) expression cut-off greater than 1.5 (Appendix Dataset S3). Gene expression was quantified using BEDTools multicov and normalised to RPKM. Gene ontology enrichment was performed using ShinyGO (Ge *et al*, 2020).

### Stress treatment and phenotyping

For all the analyses and phenotyping at least 30 plants were used. Seeds obtained from genotype-confirmed T2 plants were sterilised using ethanol, bleach and 0.1% HgCl_2_ and germinated on ½ MS media with 0.3% Phytagel (Sigma Aldrich) and for salt stress 120 mM NaCl was supplemented. Seedlings were germinated in dark for 5 days and then grown in light for further 14 days before phenotyping.

### NCPs stability assay

NCP stability assay using SYPRO orange dye was performed as described earlier (Osakabe *et al*, 2018). Fluorescence was measured on a BioRad CFX96 real time system in a FRET mode following manufacturer’s protocol and the dF/dT computation was normalised to base fluorescence at 26 ᵒC. For gel mobility shift assays, 100 ng (DNA mass) of NCPs in a 15 µl NCP buffer was treated for 1 min at mentioned temperatures in a dry bath and immediately placed back in ice. Post treatment, the NCPs mixed with native loading dye was electrophoresed on a 6% acrylamide gel in 0.5x TBE buffer.

### Immunoblotting

Total nuclear protein from 0.4 g rice seedlings were extracted as described earlier (Lorković *et al*, 2017). The proteins were electrophoresed on a 14% SDS-PAGE gel and blotted onto Protran supported nitrocellulose membrane as described (Nair *et al*, 2023). The membrane was hybridised with α-H4K5Ac (Merck 07-327, 1:2500), α-H3 (Merck 07-10254, 1:20000), α-H4 (Abcam ab10158, 1:2500) or α-H4.V (Custom generated, 1:1000) in a 5% milk containing 1x TBST buffer. For the bacterial proteins, 25 µl culture of 2.0 OD induced cells were lysed in equal volume 2x Laemmli buffer.

### Multi-stress transcriptome dataset analyses

Gene expression datasets were downloaded from TENOR (Kawahara *et al*, 2016) website (https://tenor.dna.affrc.go.jp/). Fold changes in seedling-shoot gene expression was calculated for all kinds of stresses by normalising to the corresponding control dataset. The fold change matrix was subset for the DEGs of interest, merged with the similarly processed *h4.v KO* and salt stress datasets and the non-zero entries were taken for principal component analyses using prcomp package in R. The data visualisation was done using the tools of factoextra package.

## Data availability

All raw and processed sequencing data generated in this study have been submitted to the NCBI Gene Expression Omnibus (GEO; https://www.ncbi.nlm.nih.gov/geo/) under accession number XXXXX. Coordinates and the cryo-EM maps for the rice H4 NCP and H4.V_S_ NCP have been deposited in the Protein Data Bank (PDB) and Electron Microscopy Data Bank (EMDB). Accession codes are: H4 NCP (PDB:8Q15, EMDB: EMD-18060) and H4.V_S_ NCP (PDB:8Q16, EMDB: EMD-18061).

## Acknowledgements

We thank Prof. K. Veluthambi for *Agrobacterium* strains, rice seeds, and binary plasmids. We thank NGGF, CIFF, IT, radiation, cryo-EM, greenhouse and laboratory-kitchen facilities at the NCBS. We would like to thank Dr Katja Lammens for the technical help during cryo-EM data collection and Prof. Dr. Karl-Peter Hopfner and team as well as Dr. Ramesh Yelagandula for useful discussions.

## Author Contributions

PVS and VHS designed the study. PVS obtained funding, analysed the data and wrote the manuscript along with VHS. VHS performed all the genetic, genomic, biochemical and molecular biology experiments and generated and reconstituted NCPs. PSP and AAJ generated cryo-EM structures and performed its analysis. FG contributed to cryo-EM data acquisition. SR performed most of the microscopy analysis. SJ, AG and VKR optimized cryo-EM. FB provided reagents and guidance on specific aspects. All the authors have read and approved the manuscript.

## Appendix Information

**Appendix Figure S1.**
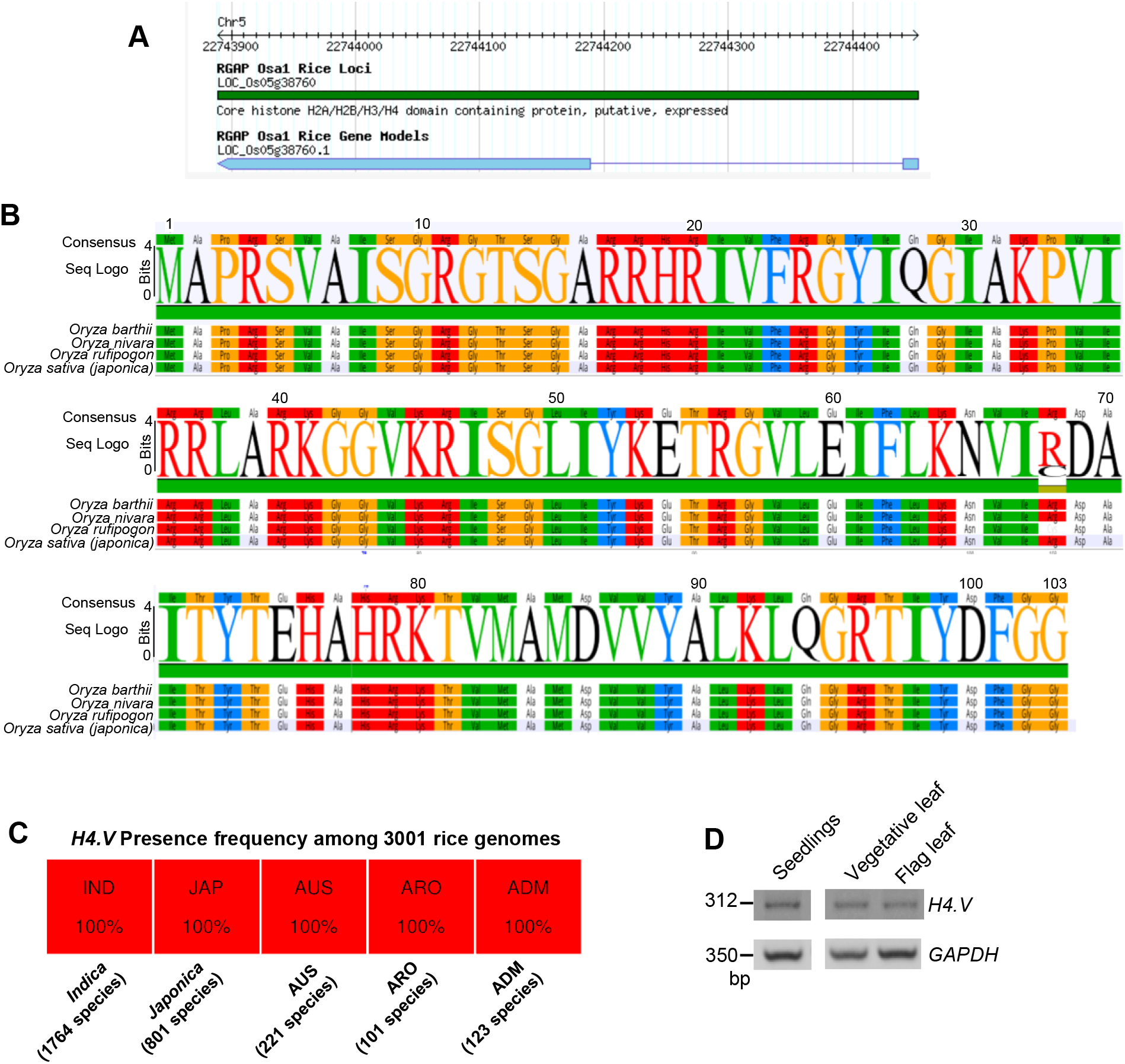
H4.V is conserved among Oryza genera. (A) Gene model of the H4.V from RAPDB (https://rapdb.dna.affrc.go.jp/). (B) Seq-logo showing conservation of the H4.V among rice species. (C) H4.V presence frequency among the 3001-rice genome project from rice pan-genome browser (https://cgm.sjtu.edu.cn/3kricedb/index.php). (D) RT-PCR amplification of the H4.V transcript across three rice tissues. GAPDH served as control.

**Appendix Figure S2.**
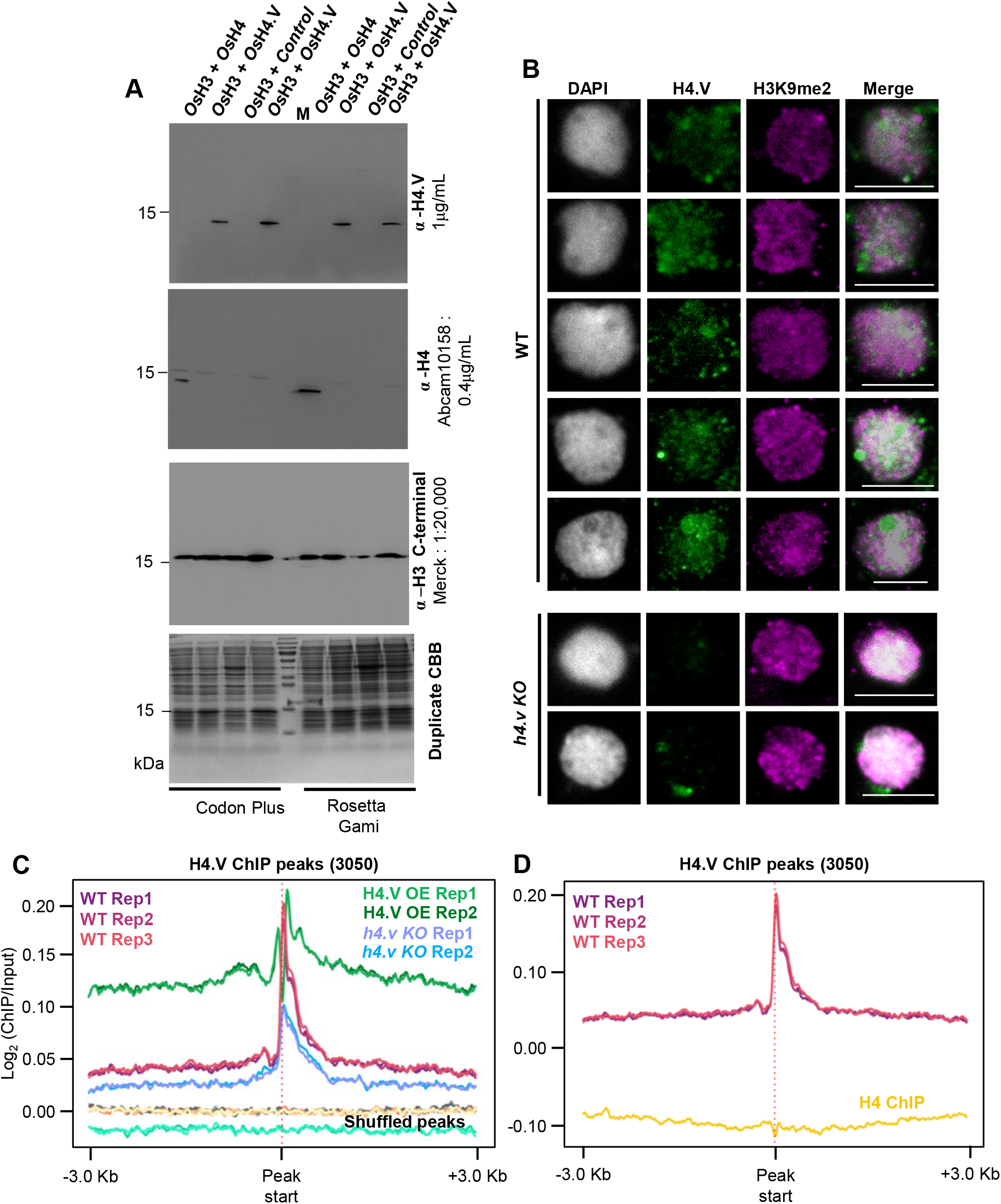
H4.V antibody is specific and does not cross react with H4. (A) Immunoblots showing reactivity of α-H4.V and α-H4 specifically to their epitopes. Total bacterial lysates from two different *E. coli* strains are shown. Blots were stripped and re-hybridised. CBB stained duplicate gel served as loading control. Both H3 and H4/H4.V were co-expressed in the same plasmid (Appendix Table S5). M: size marker. (B) IFL images of nuclei from WT and *h4.v KO* plants stained using α–H4.V and α–H3K9me2. Scale: 5 µm. (C) ChIP enrichment profiles of H4.V at H4.V peaks from WT, OE-H4.V and *h4.v KO*. Enrichment profiles at shuffled H4.V peaks served as control. (D) ChIP enrichment profiles at H4.V peaks from H4.V ChIP-seq and H4 ChIP-seq.

**Appendix Figure S3.**
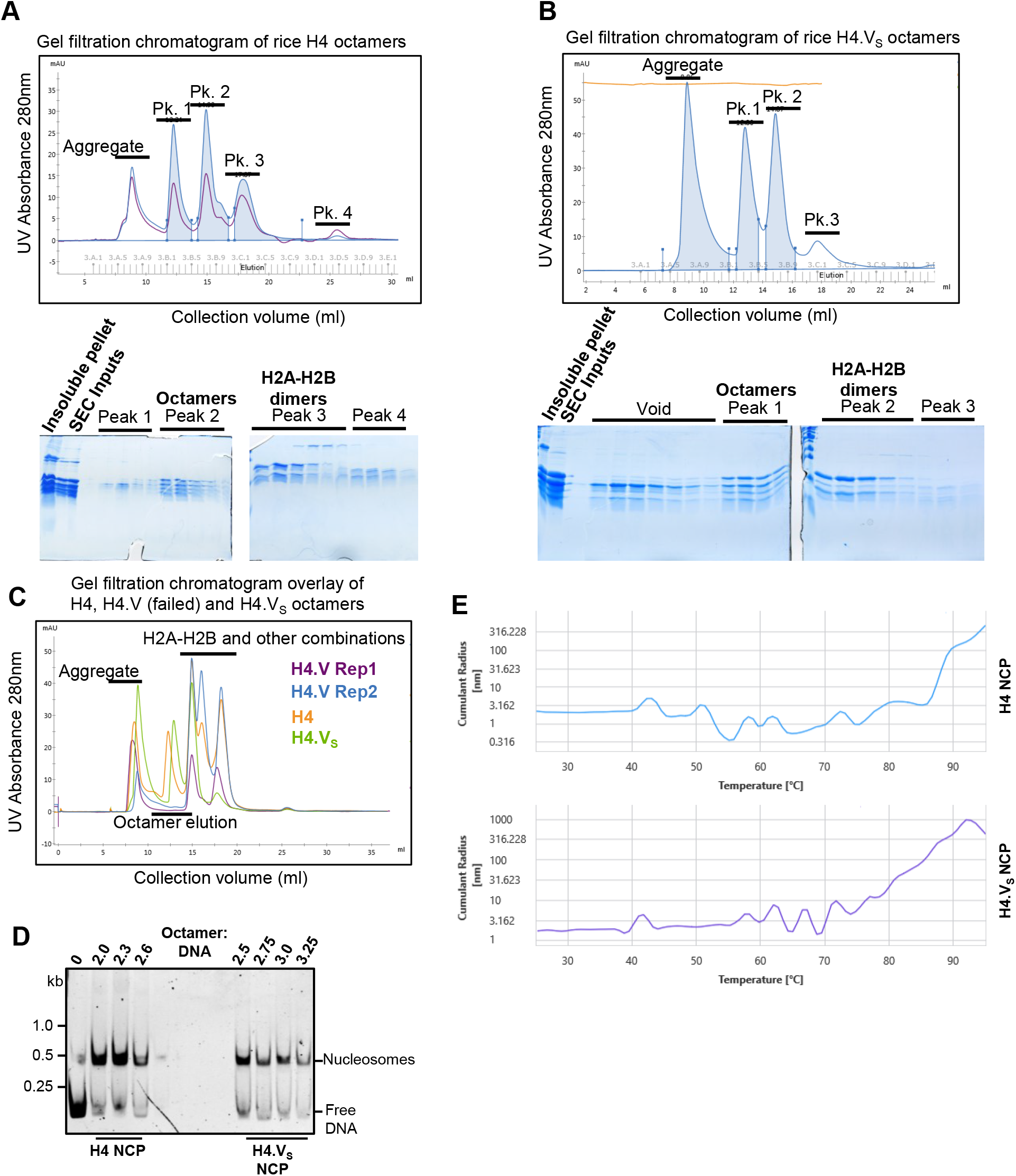
H4.V can form nucleosomes in vitro. (A-B) Gel filtration chromatograms of the serially dialysed histone refolding mix containing H4 (A) or H4.V_S_. (B) with representative fractions of peaks resolved on CBB stained SDS-PAGE gels. Portion of the chromatograms are (C) Overlay of gel filtration chromatograms of H4, H4.V_S_ with that of H4.V. (D) Gel-mobility shift assays showing titration of the NCPs with different molar combinations of octamer and Widom 601 NPS (146 bp DNA). (E) DLS plots of estimated hydrodynamic sizes of the NCPs over the temperature range.

**Appendix Figure S4.**
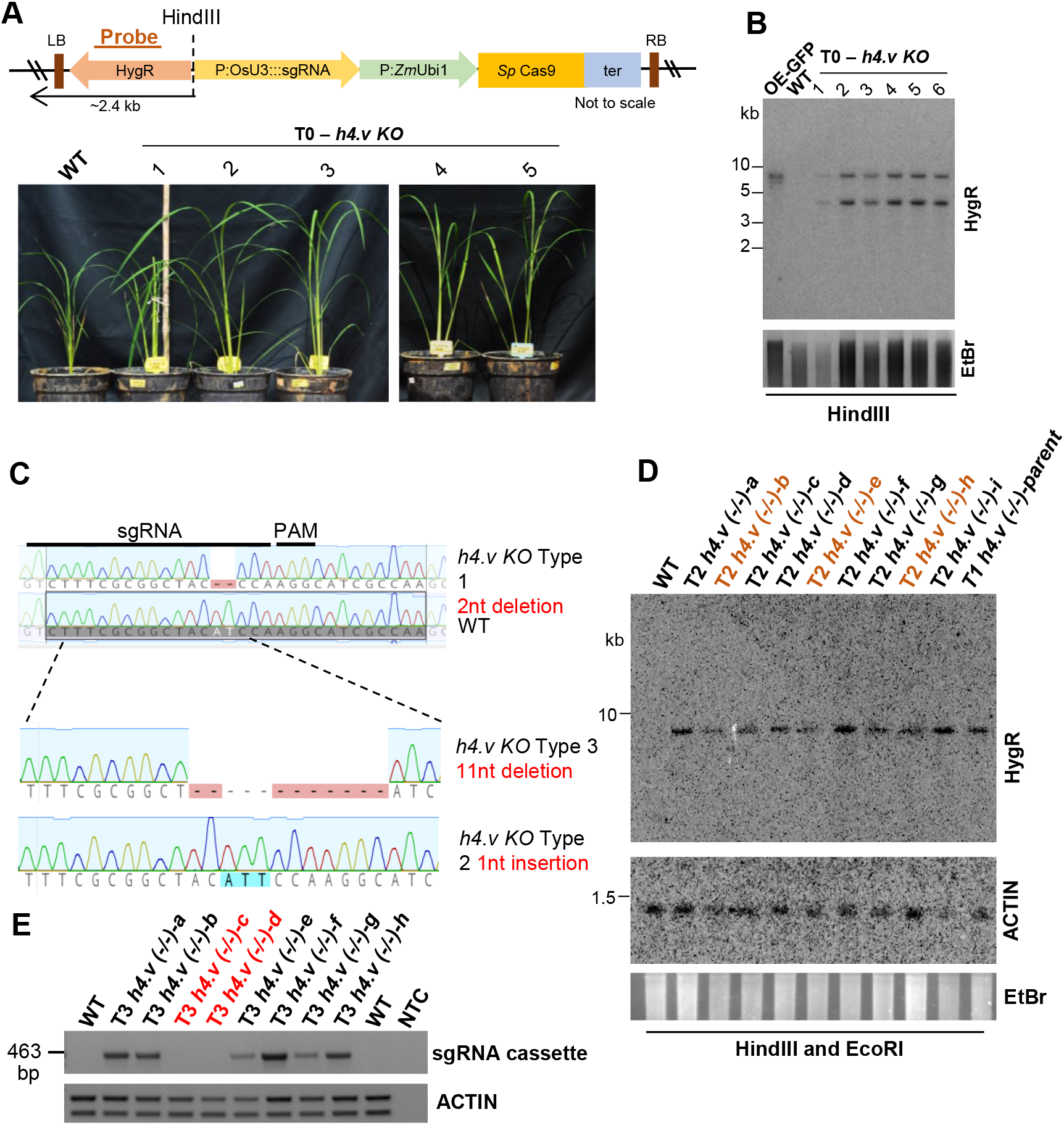
Generation of CRISPR-Cas9 mediated T-DNA free *h4.v KO* in rice. (A) T-DNA map of the Cas9 construct used for generating the *h4.v KO*. T0 *h4.v KO* images of 5-weeks old plants are shown. (B) Southern blots showing the junction fragment profiles of T0 *h4.v KO* lines. OE-GFP served as positive control. (C) Mutation profiles of T1 *h4.v KO* plants compared to WT. Three types of mutations were obtained and the type 1 mutation was taken for the analysis. (D) Junction fragment Southern analysis of the homozygous (-/-) T1 *h4.v KO* parent (that segregated as single copy transgene) and its T2 progeny. Plants marked in brown are putative parents that are heterozygous for the transgene. (E) Genomic DNA PCR showing the T3 segregants of the T2 parents (marked brown in (D)). Actin was used a loading control and sgRNA region as transgene marker. (A, B and D) Probe (HygR) used for T-DNA junction fragment Southern blots is marked (brown). EtBr-stained gel served as loading control.

**Appendix Figure S5.**
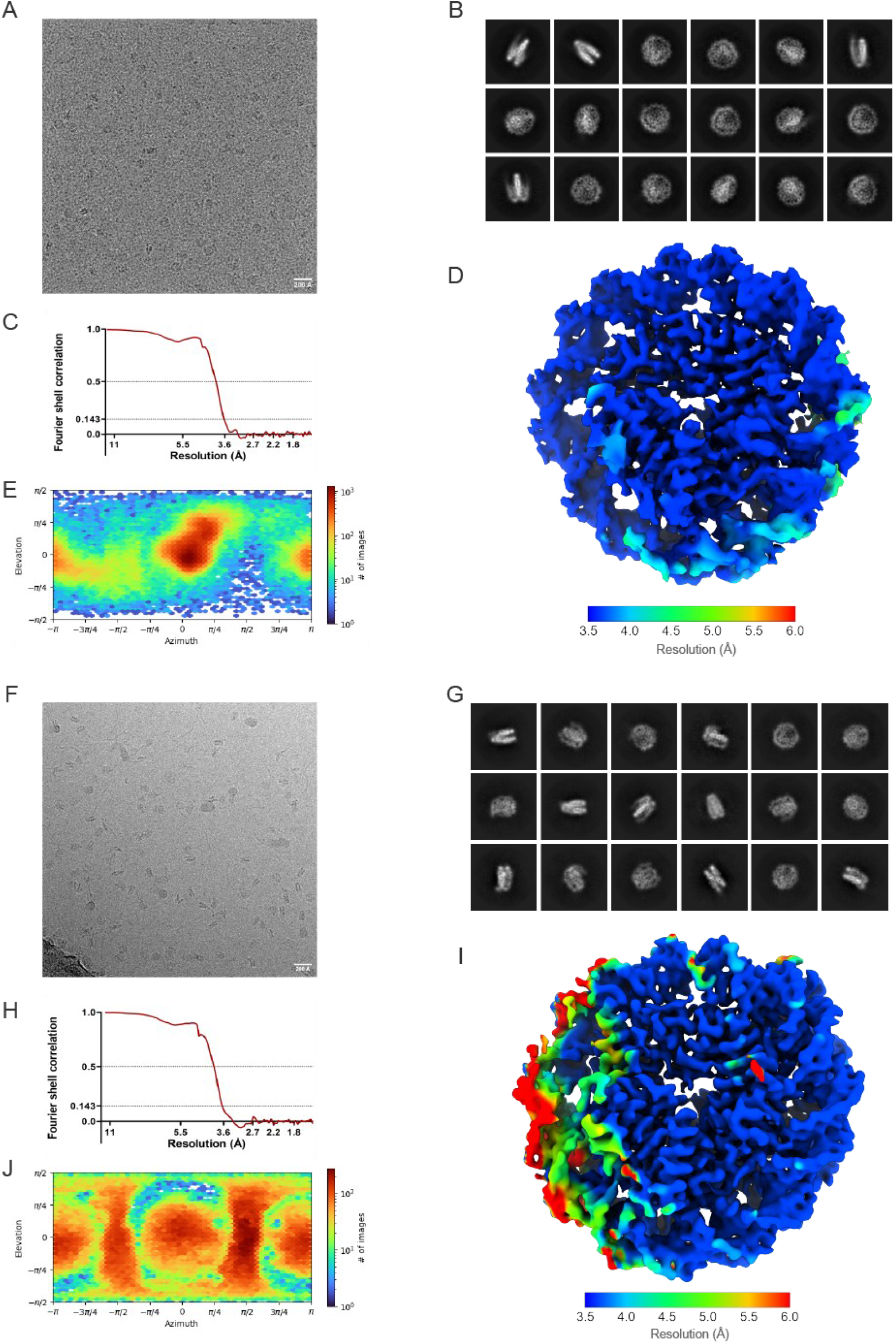
Cryo-EM data collection and image processing of rice NCPs. (A-J) Data collection and refinement for rice canonical NCP (A-E) and H4.Vs NCP (F-J). Representative cryo-EM micrographs (A,F), representative 2D classes (B,G), Fourier Shell Correlation curves (C,H) showing the resolution estimation of the maps, local resolution maps (D,J) and angular distribution of the particles (E,J) for the NCPs were depicted.

**Appendix Figure S6.**
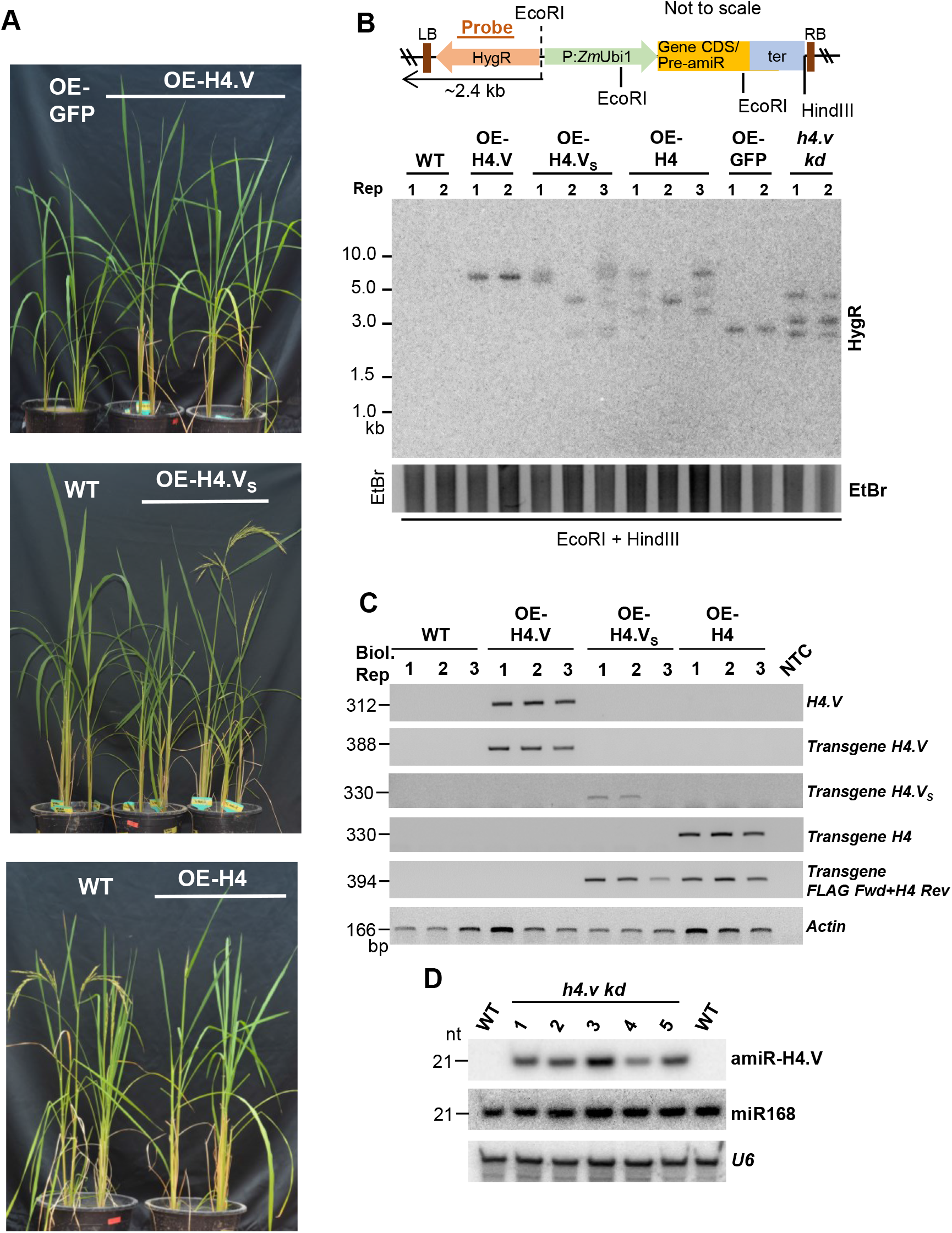
Phenotypes and Southern blot validation of H4.V expression perturbed plants. (A) Phenotypes of plants overexpressing H4.V, H4.V_S_ and H4. OE-GFP served as vector control. (B) T-DNA map and the junction fragment (arrow) Southern analyses of transgenic plants overexpressing H4.V, H4.V_S_ and H4 or GFP (3x-FLAG tagged at the N-terminal) or the precursor of amiR. HygR was used as probe (brown bar). EtBr-stained gel is the loading control. (C) RT-PCR (semi-quantitative) gels showing the OE of the transgenes in leaves using specific primers. NTC-no template control. Actin was used as loading control. (D) sRNA northern blots showing the accumulation of the amiR targeting H4.V in *h4.v kd* plants. U6 served as control.

**Appendix Figure S7.**
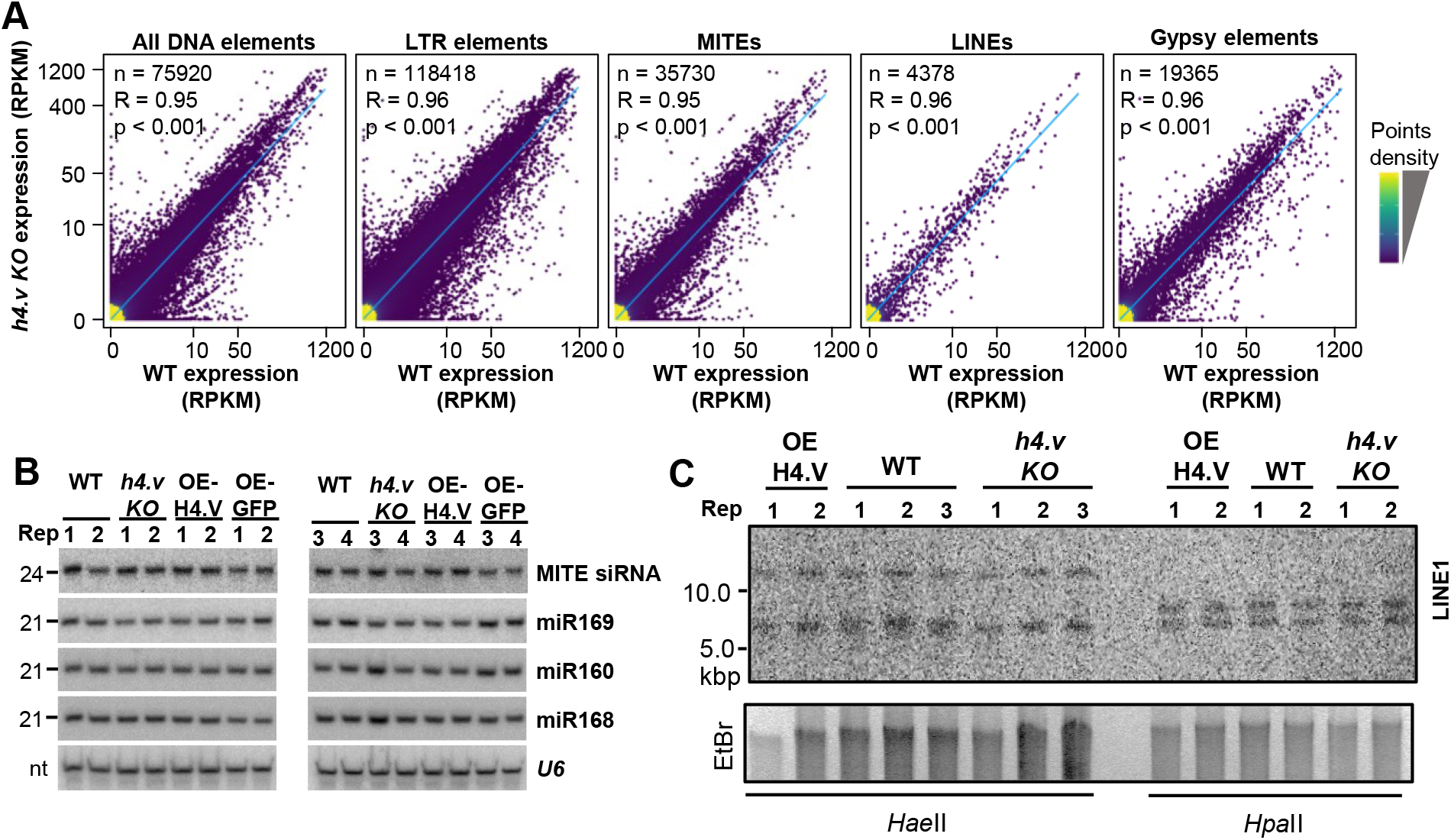
Transposon regulation is unaffected in *h4.v KO* plants. (A) Density scatter plots showing expression difference of different categories of transposons. Pearson correlation coefficient (R) and p-values are mentioned. Points density are mentioned with colour gradient. (B) sRNA northern blots showing abundance of repeat derived sRNAs or other miRNAs. U6 was the loading control. (C) Methylation sensitive Southern blot hybridised with LINE1 probe. EtBr-stained gel served as loading control.

**Appendix Figure S8.**
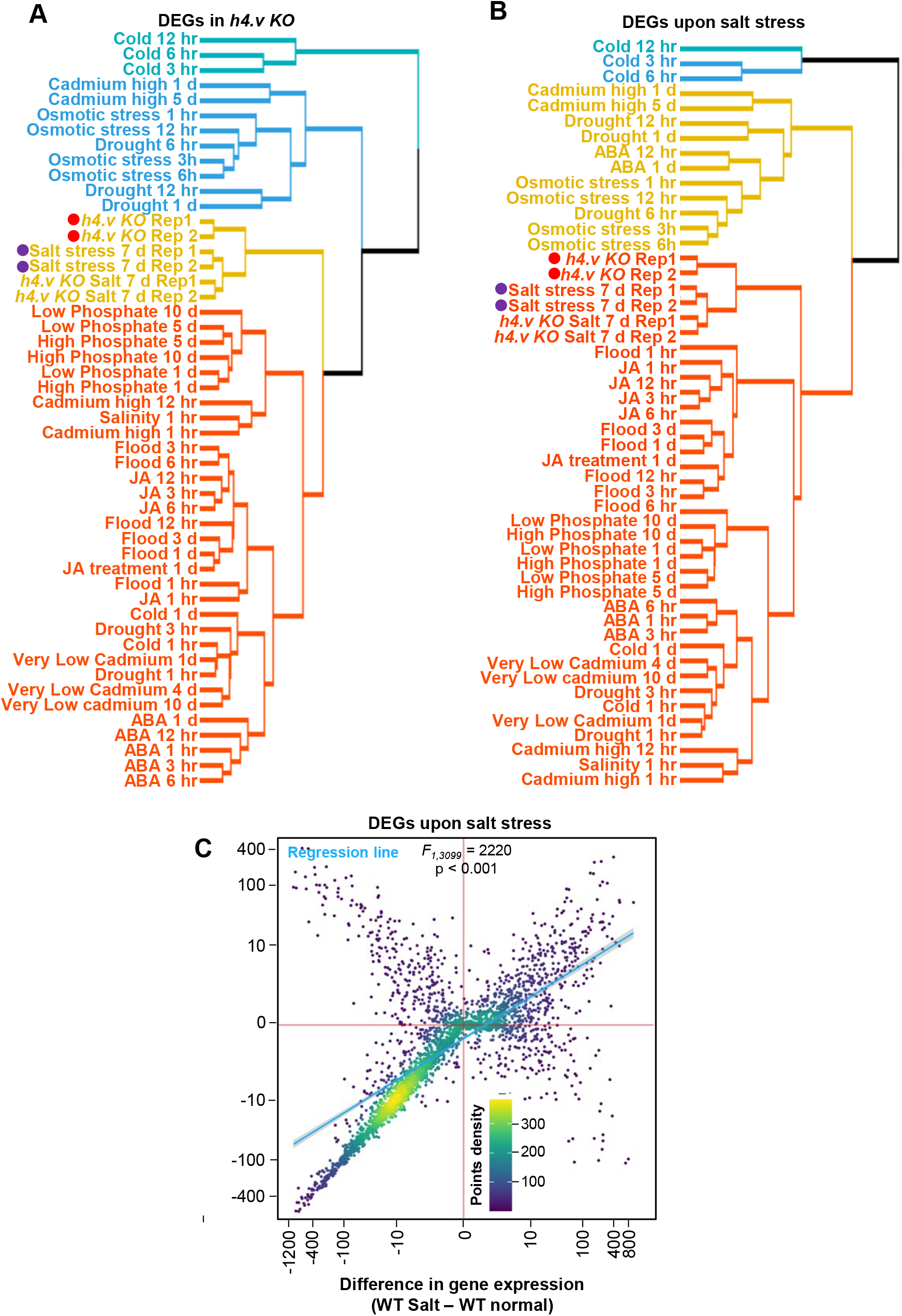
H4.V is necessary to prevent precocious salt stress like transcriptome. (A-B) Clustered dendrograms comparing transcriptomes across different stress conditions from TENOR datasets, analysed for the *h4.v KO* DEGs (A), or the salt stress DEGs (B). Red dots highlight the *h4.v KO* profiles and purple represent salt stress profiles. Fig 5C and D depict representative examples from stress types and replicates are shown here. The tree is clustered into four hierarchical types. (C) Density scatter plots showing resemblance of *h4.v KO* with salt stress transcriptomes for the salt stress specific DEGs. Details are in Fig. 5G.

**Appendix Figure S9.**
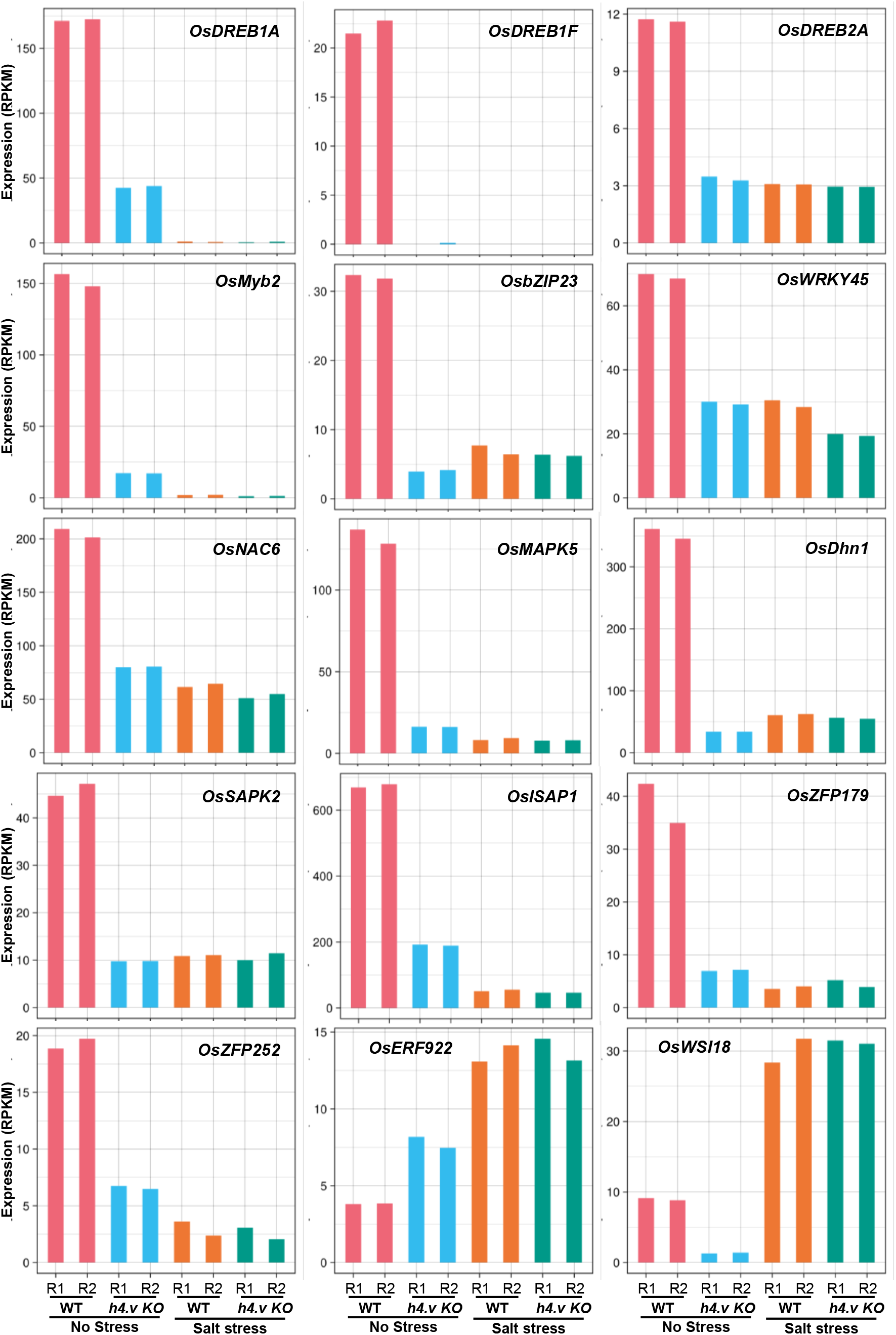
Expression of salt-stress responsive genes. Gene expression levels in WT and *h4.v KO* with and without salt stress. Gene names are mentioned.

**Figure S10.**
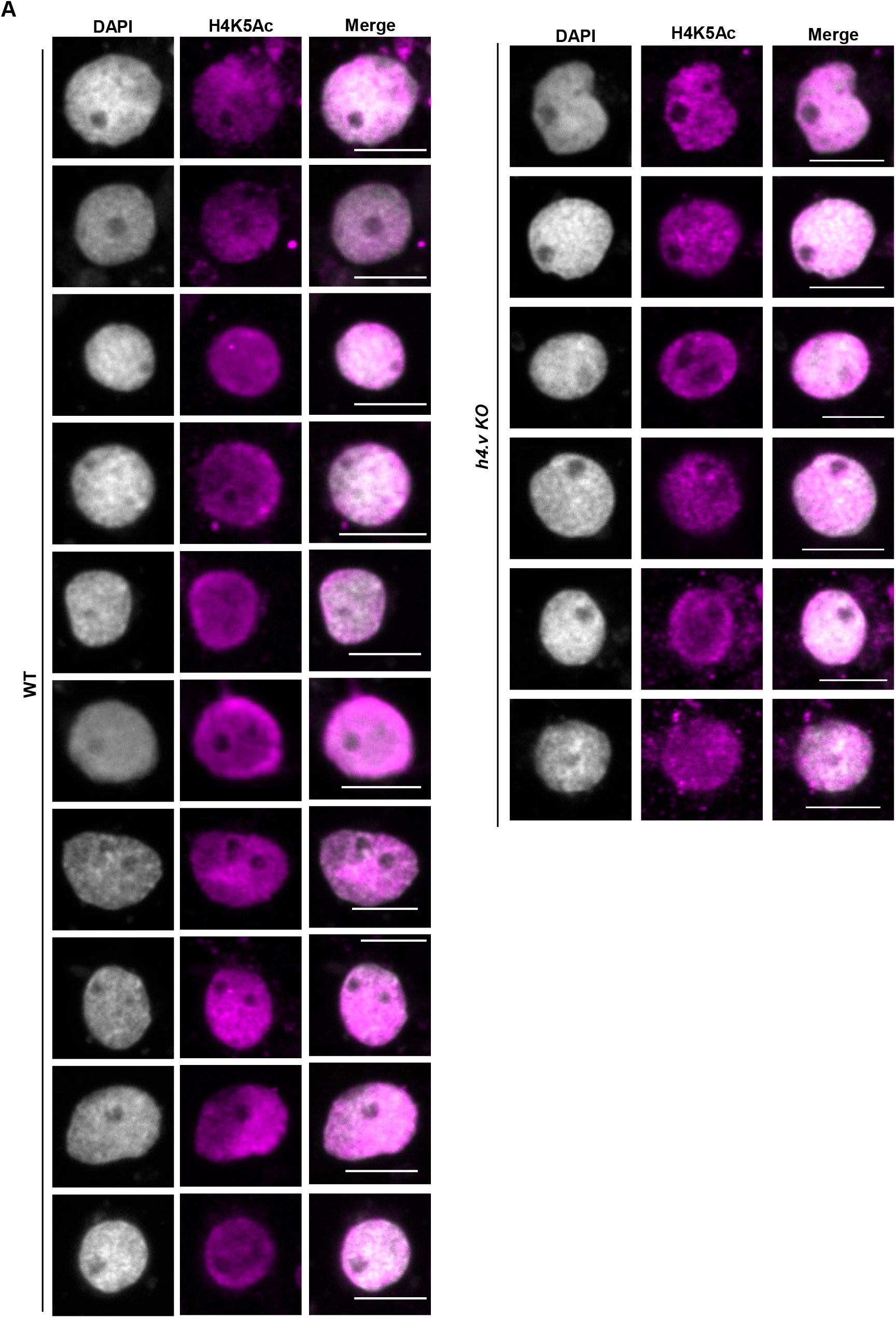
Perturbation of H4 variant does not cause global decline of H4K5Ac marks. (A) IFL images showing occupancy of H4K5Ac marks in WT and *h4.v KO*.

---

**Appendix Table S1.**
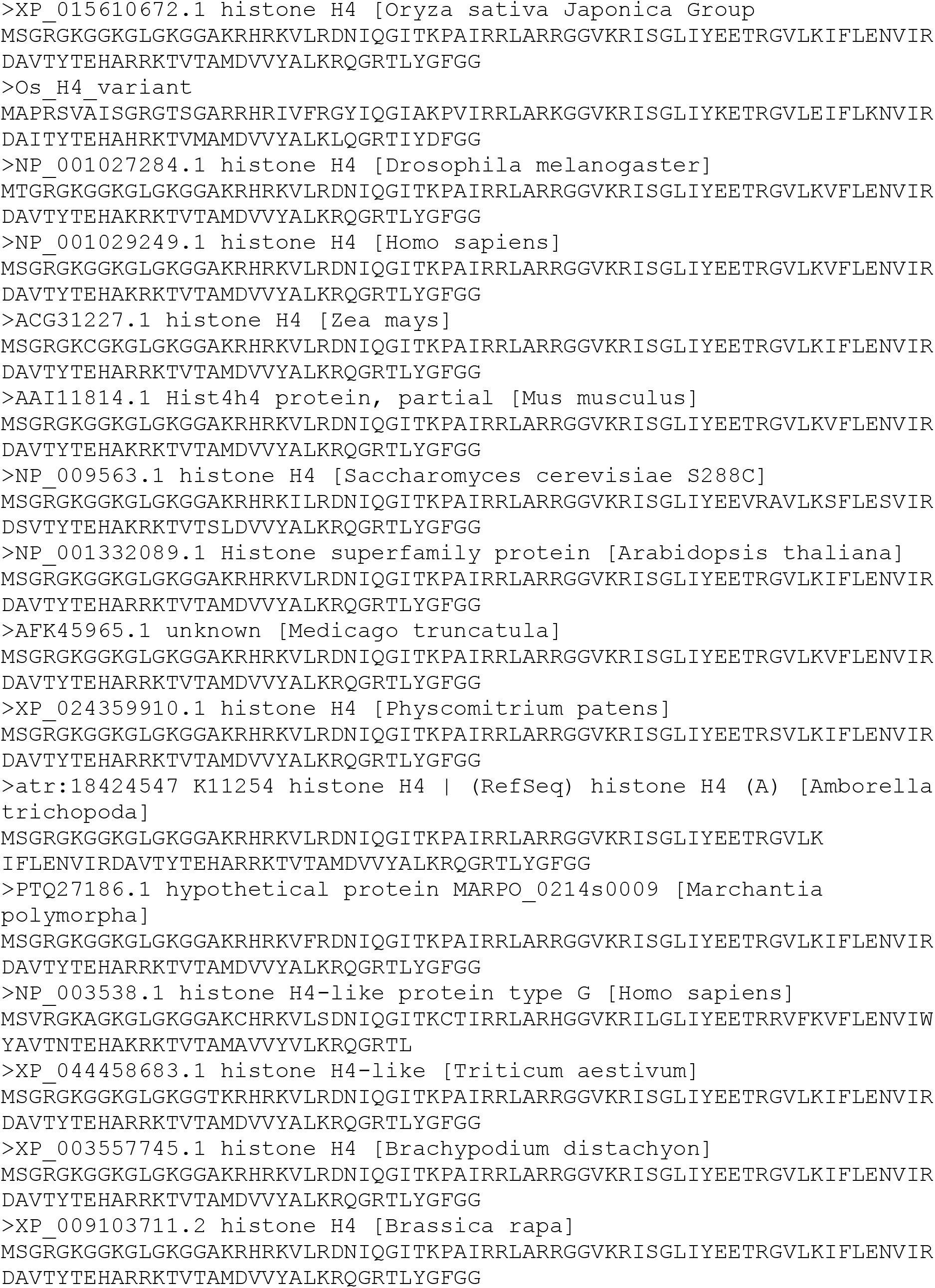
Histone H4 sequences from different species.

**Appendix Table S2.** Details of high-throughput genomics data generated in this study.

| Sl. No. | Dataset type | Genotype | Replicate | Source tissue | Sequencing mode |
| --- | --- | --- | --- | --- | --- |
| 1 | RNA-seq | WT | Rep1 | Seedlings without stress. 14 days old. | Paired end-100bp |
| 2 | RNA-seq | WT | Rep2 |  | Paired end-100bp |
| 3 | RNA-seq | *h4.v KO* | Rep1 |  | Paired end-100bp |
| 4 | RNA-seq | *h4.v KO* | Rep2 |  | Paired end-100bp |
| 5 | RNA-seq | WT | Rep1 | Seedlings with 120 mM salt stress. 14 days old. | Paired end-100bp |
| 6 | RNA-seq | WT | Rep2 |  | Paired end-100bp |
| 7 | RNA-seq | *h4.v KO* | Rep1 |  | Paired end-100bp |
| 8 | RNA-seq | *h4.v KO* | Rep2 |  | Paired end-100bp |
| 9 | RNA-seq | WT | Rep1 | Mature Leaves from 60 days old plants. | Paired end-100bp |
| 10 | RNA-seq | WT | Rep2 |  | Paired end-100bp |
| 11 | RNA-seq | OE-H4.V | Rep1 |  | Paired end-100bp |
| 12 | RNA-seq | OE-H4.V | Rep2 |  | Paired end-100bp |
| 13 | RNA-seq | OE-H4.V<sub>s</sub> | Rep1 |  | Paired end-100bp |
| 14 | RNA-seq | OE-H4.V<sub>s</sub> | Rep2 |  | Paired end-100bp |
| 15 | RNA-seq | OE-H4 | Rep1 |  | Paired end-100bp |
| 16 | RNA-seq | OE-H4 | Rep2 |  | Paired end-100bp |
| 17 | ChIP-seq with α-H4.V | WT | Rep1 | Seedlings without stress. 14 days old. | Single end-50bp |
| 18 | ChIP-seq with α-H4.V | WT | Rep2 | | Single end-50bp |
| 19 | ChIP-seq with α-H4.V | WT | Rep3 | | Single end-50bp |
| 20 | ChIP-seq with α-H4.V | *h4.v KO* | Rep1 | | Single end-50bp |
| 21 | ChIP-seq with α-H4.V | *h4.v KO* | Rep2 | | Single end-50bp |
| 22 | ChIP-seq with α-H4.V | WT | Rep1 | Seedlings with 120 mM salt stress. 14 days old. | Single end-50bp |
| 23 | ChIP-seq with α-H4.V | WT | Rep2 | | Single end-50bp |
| 24 | ChIP-seq with α-H4K5Ac | WT | Rep1 | Seedlings without stress. 14 days old. | Single end-50bp |
| 25 | ChIP-seq with α-H4K5Ac | WT | Rep2 | | Single end-50bp |
| 26 | ChIP-seq with α-H4K5Ac | *h4.v KO* | Rep1 | | Single end-50bp |
| 27 | ChIP-seq with α-H4K5Ac | *h4.v KO* | Rep2 | | Single end-50bp |
| 28 | ChIP-seq with α-H4K5Ac | WT | Rep1 | Seedlings with 120 mM salt stress. 14 days old. | Single end-50bp |
| 29 | ChIP-seq with α-H4K5Ac | WT | Rep2 | | Single end-50bp |
| 30 | ChIP-seq with α-H4K5Ac | *h4.v KO* | Rep1 | | Single end-50bp |
| 31 | ChIP-seq with α-H4K5Ac | *h4.v KO* | Rep2 | | Single end-50bp |
| 32 | ChIP-seq with α-H4K5Ac | OE-H4.V | Rep1 | Seedlings without stress. 14 days old. | Single end-50bp |
| 33 | ChIP-seq with α-H4K5Ac | OE-H4.V | Rep2 | | Single end-50bp |
| 34 | ChIP-seq with α-H4.V | OE-H4.V | Rep1 | | |
| 35 | ChIP-seq with α-H4.V | OE-H4.V | Rep2 | | |
| 36 | ChIP-seq with α-H4C | WT | Rep1 | | Single end-50bp |
| 37 | ChIP-seq with α-H4C | *h4.v KO* | Rep1 | | Single end-50bp |
| 38 | MNase sensitivity on NCPs | Undigested Input DNA | Rep1 | Synthetic NCPs | Paired end-100bp |
| 39 | MNase sensitivity on NCPs | H4 NCPs | Rep1 | Synthetic NCPs | Paired end-100bp |
| 40 | MNase sensitivity on NCPs | H4.V NCPs | Rep1 | Synthetic NCPs | Paired end-100bp |

**Appendix Table S3.** Details of high-throughput genomics data obtained from publicly available datasets.

| Sl. No. | Dataset type | Genotype | Source tissue | SRA number | GSE number | Reference |
| --- | --- | --- | --- | --- | --- | --- |
| 1 | ChIP-seq with α-H3K9me2 Rep1 | WT | seedlings | SRX7426641 | GSE142462 | (Zhao *et al*, 2020) |
| 2 | ChIP-seq with α-H3K9me2 Rep2 | WT | seedlings | SRX7426642 | GSE142462 | (Zhao *et al*, 2020) |
| 3 | ChIP-seq with α-H2A.Z Rep1 | WT | Seedlings | SRX2860628 | GSE155269 | (Du *et al*, 2020) |
| 4 | ChIP-seq with α-H2A.Z Rep2 | WT | Seedlings | SRX2860589 | GSE155269 | (Du *et al*, 2020) |
| 5 | ChIP-seq with α-H2A.Z Rep3 | WT | Seedlings | SRX1883882 | GSE155269 | (Du *et al*, 2020) |
| 6 | ChIP-seq with α-H3K27me3 Rep1 | WT | Seedlings | SRX1131812 | GSE71640 | (Zhou *et al*, 2016) |
| 7 | ChIP-seq with α-H3K27me3 Rep2 | WT | Seedlings | SRX2004187 | GSE71640 | (Zhou *et al*, 2016) |
| 8 | ChIP-seq with α-H4K12Ac Rep1 | WT | Seedlings | SRX038893 | GSE26734 | (Zhang *et al*, 2012) |
| 9 | ChIP-seq with α-H4K16Ac Rep1 | WT | Seedlings | SRX1044780 | GSE69426 | (Lu *et al*, 2015) |
| 10 | Input_ChIP-seq_Rep1 | WT | Seedlings | SRX10085735 | GSE166665 | (Pachamuthu *et al*, 2022) |
| 11 | Input_ChIP-seq_Rep2 | WT | Seedlings | SRX10085736 | GSE166665 | (Pachamuthu *et al*, 2022) |

**Appendix Table S4.**
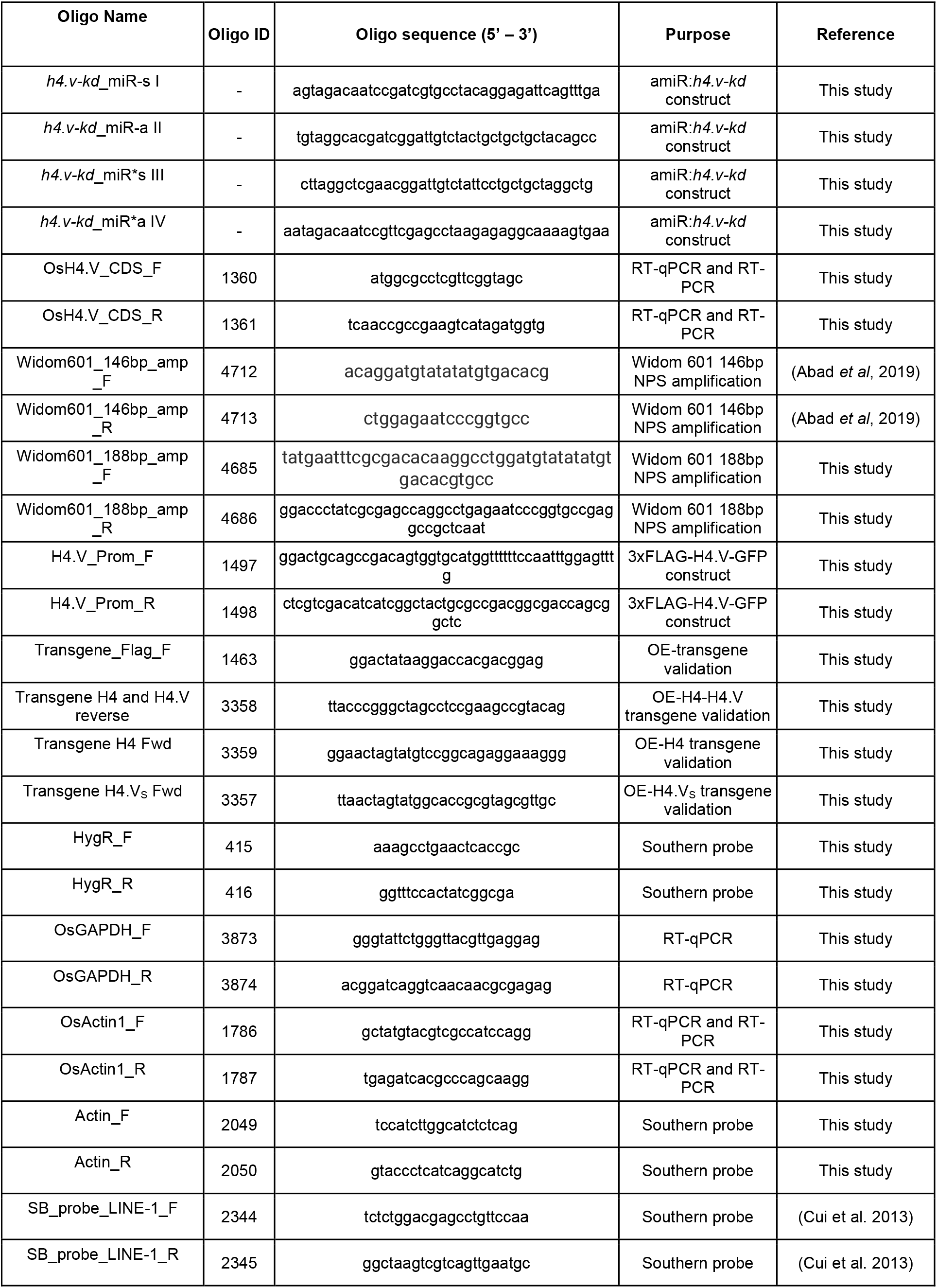
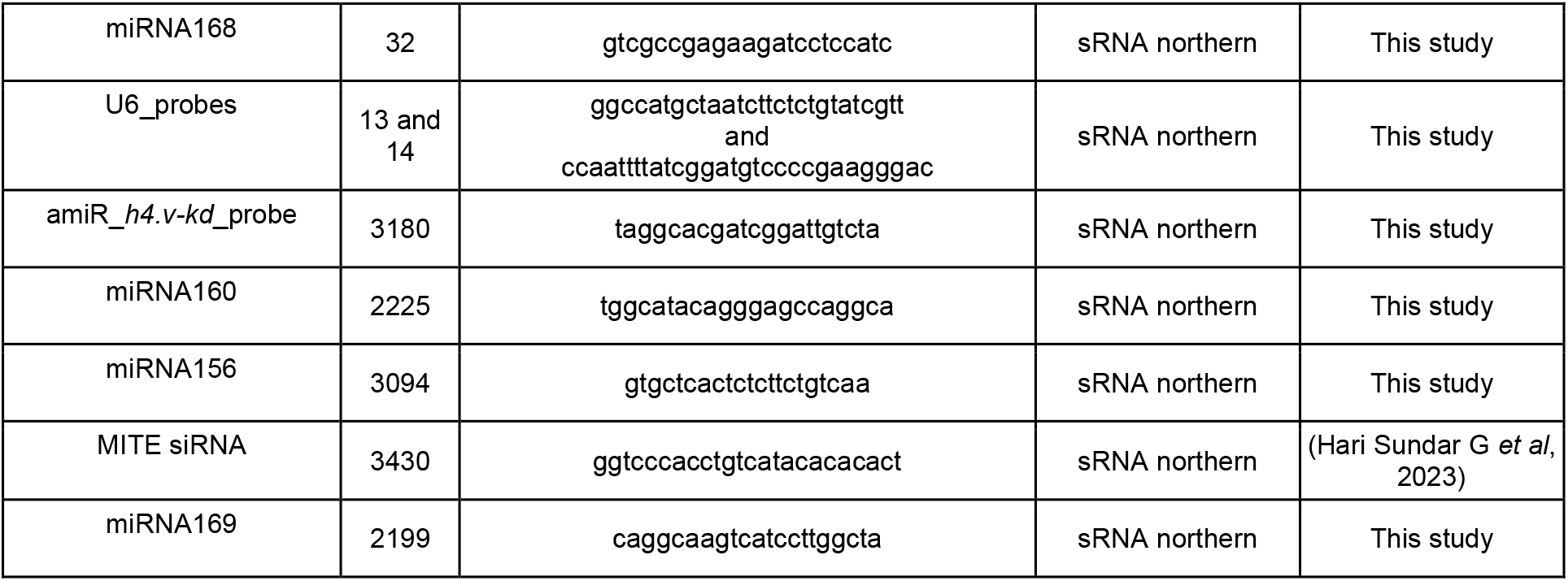
List of oligos and probes used in this study.

**Appendix Table S5.**
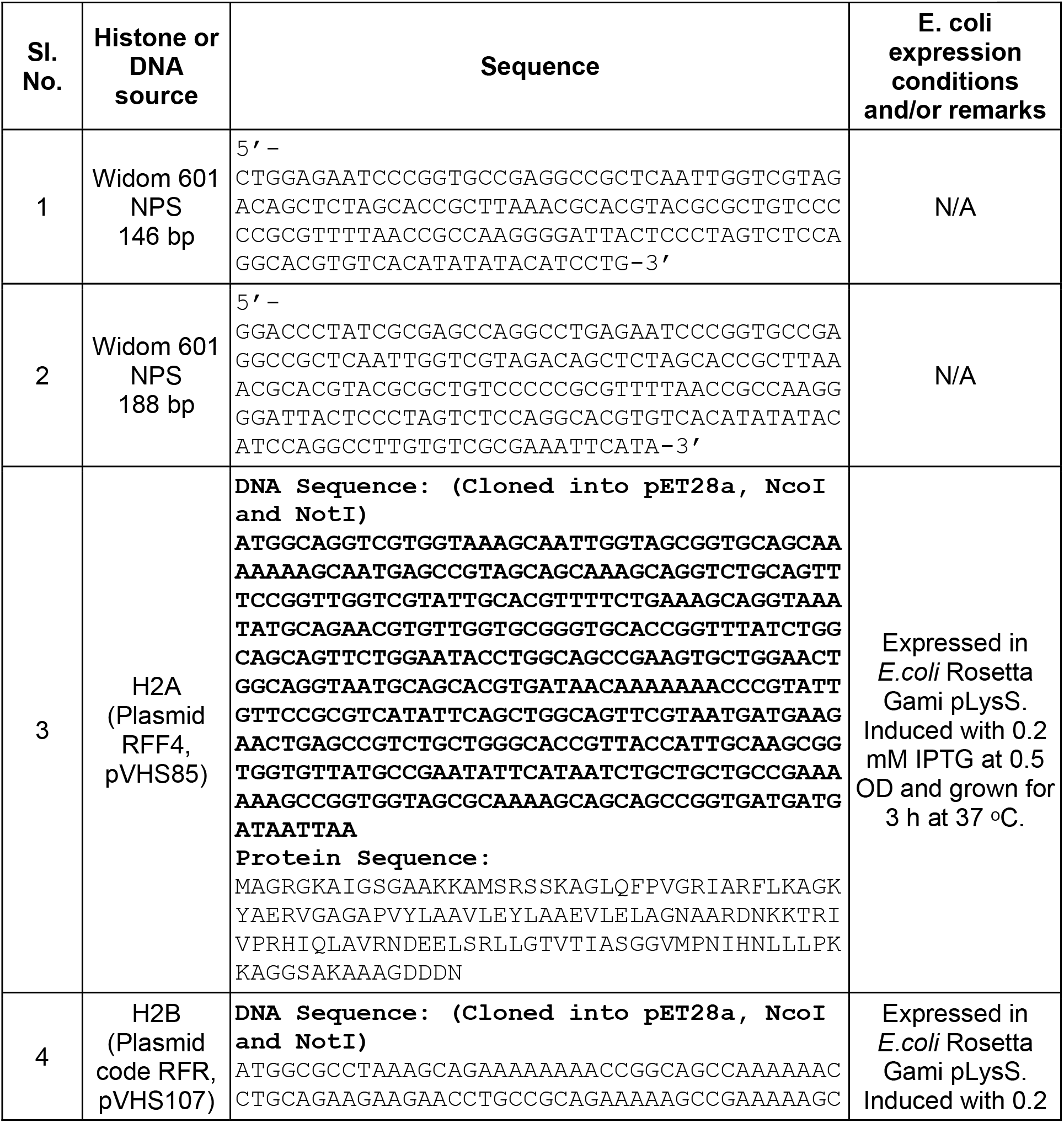
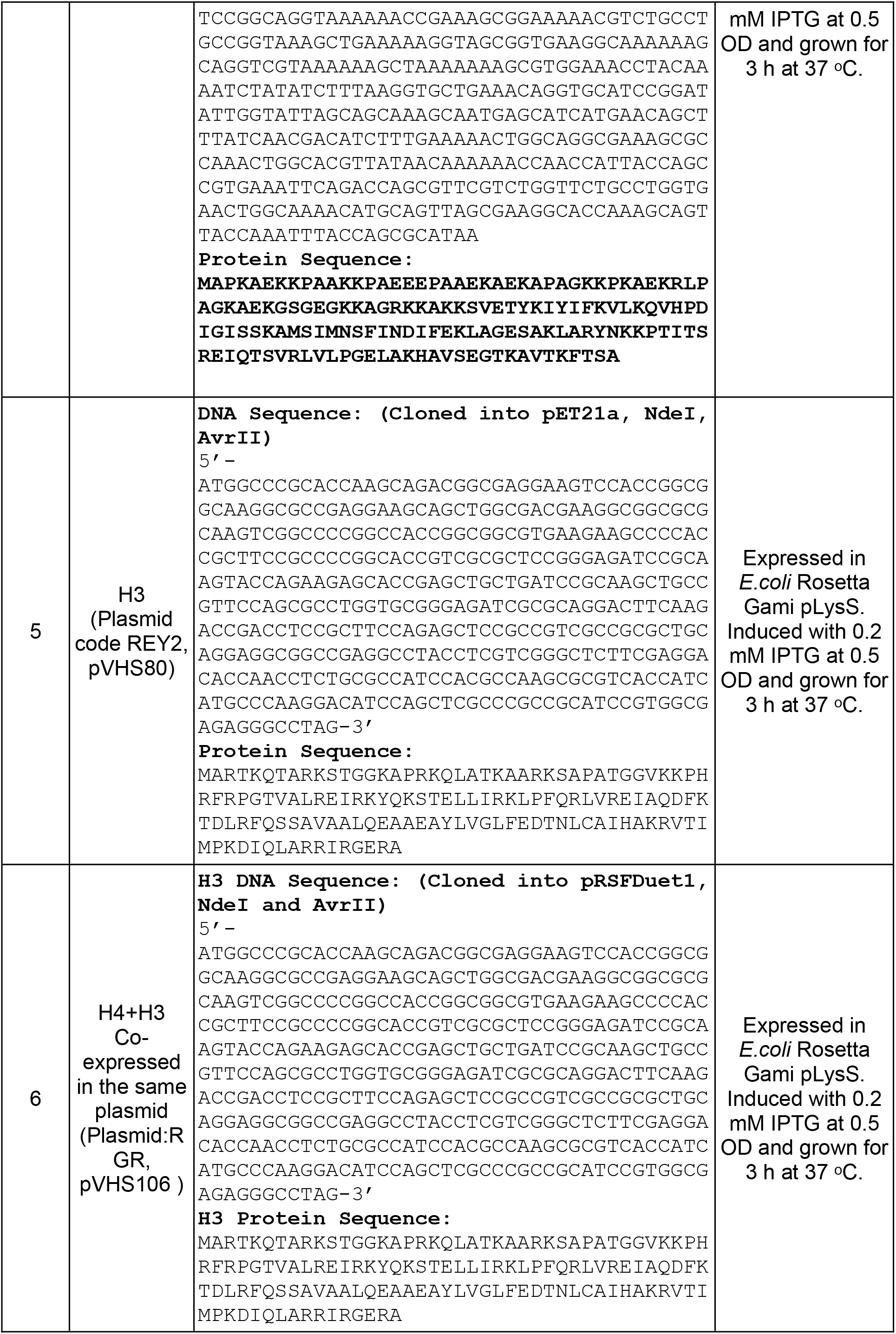
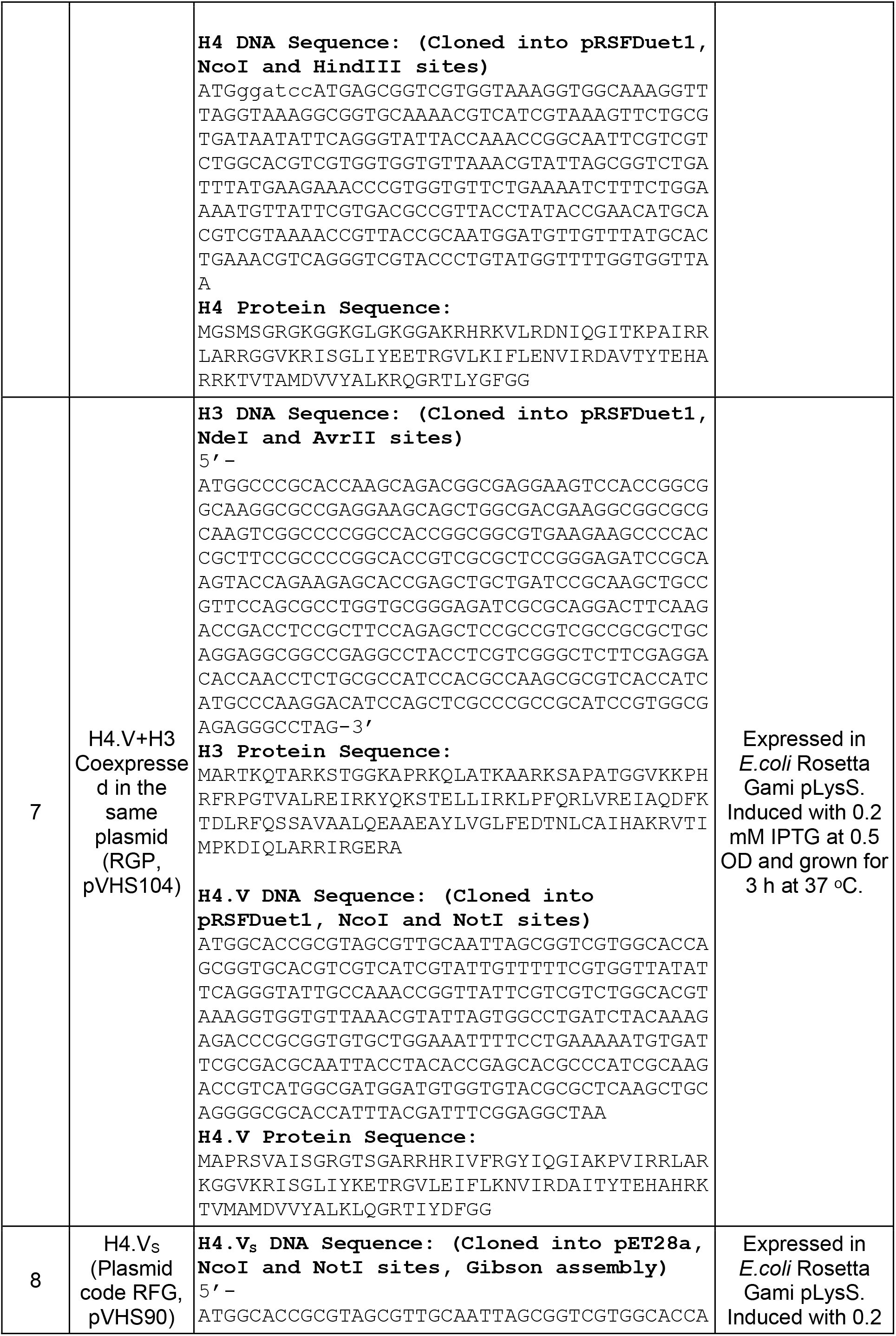
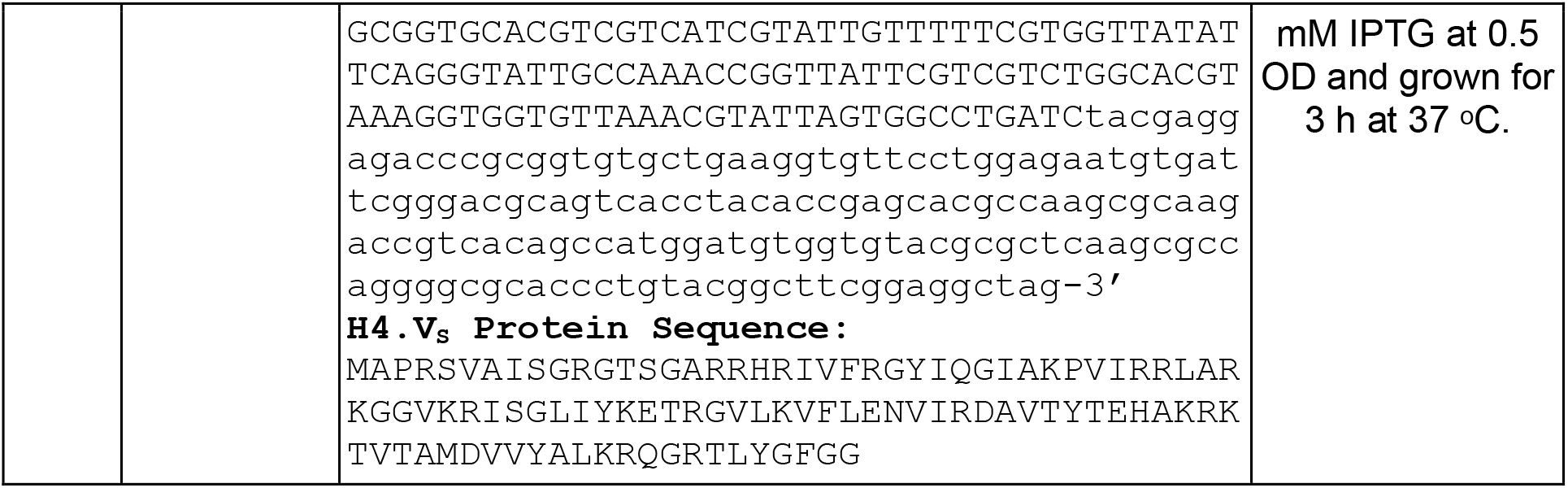
Details of *in vitro* reconstitution of NCPs using rice histones.

**Appendix Table S6:** Details of cryo-EM data collection and processing.

|  | H4 NCP | H4.Vs NCP |
| --- | --- | --- |
| Number of grids used | 1 | 1 |
| Grid type | Quantifoil R2/1 200 mesh + 2 nm carbon | Quantifoil R2/1 200 mesh |
| Microscope/detector | Titan Krios / Falcon4i | Titan Krios / Falcon4i |
| Voltage | 300 kV | 300 kV |
| Magnification | 165k | 165k |
| Recording mode | Counting mode | Counting mode |
| Dose rate | 1 e<sup>-</sup>/ Å<sup>2</sup>/frame | 1 e<sup>-</sup>/ Å<sup>2</sup>/frame |
| Defocus | -0.5 µm to -2.6 µm (step size 0.3 µm) | -0.5 µm to -2.6 µm (step size 0.3 µm) |
| Pixel size | 0.727 | 0.727 |
| Total dose | 60 e<sup>-</sup>/Å<sup>2</sup> | 60 e<sup>-</sup>/Å<sup>2</sup> |
| Number of frames/movie | 60 | 60 |
| Total exposure time | Adjusted to keep total dose stable | Adjusted to keep total dose stable |
| Number of micrographs | 10,395 | 31,174 |
| Number of micrographs used | 10,139 | 23,019 |
| Number of particles used | 140,877 | 148,217 |
| PDB | 8Q15 | 8Q16 |
| EMDB | EMD-18060 | EMD-18061 |
| Map resolution (FSC 0.143) | 3.6 | 3.6 |
| **Refinement (Phenix)** |  |  |
| Resolution (Å) | 4.0 | 3.6 |
| Map CC | 0.79 | 0.84 |
| Mean B factor (Å<sup>2</sup>) | 17.81 (protein) 88.23 (DNA) | 43.02 (protein) 129.02 (DNA) |
| **Validation** |  |  |
| All atom clashscore | 3.75 | 3.69 |
| Rotamer outliers (%) | 1.27 | 0.50 |
| MolProbity score | 1.27 | 1.32 |
| **Ramachandran plot** |  |  |
| Favored (%) | 97.86 | 97.07 |
| Outliers (%) | 0.00 | 0.00 |
| **RMS deviation** |  |  |
| Bond length (Å) | 0.003 | 0.004 |
| Bond angle (°) | 0.520 | 0.544 |

Appendix Dataset S1. Lists of ChIP peaks identified.

Appendix Dataset S2. List of DEGs identified upon salt stress and in genotypes.

